# Sox8 remodels the cranial ectoderm to generate the ear

**DOI:** 10.1101/2021.04.20.440617

**Authors:** Ailin Leticia Buzzi, Jingchen Chen, Alexandre Thiery, Julien Delile, Andrea Streit

## Abstract

The vertebrate inner ear arises from a pool of progenitors with the potential to contribute to all the sense organs and cranial ganglia in the head. Here we explore the molecular mechanisms that control ear specification from these precursors. Using a multi-omics approach combined with loss-of-function experiments we identify a core transcriptional circuit that imparts ear identity, along with the first genome-wide characterization of non-coding elements that integrate this information. This analysis places the transcription factor Sox8 at the top of the ear determination network. Introducing Sox8 into cranial ectoderm not only converts non-ear cells into ear progenitors, but also activates the cellular programs for ear morphogenesis and neurogenesis. Thus, Sox8 has the unique ability to remodel transcriptional networks in the cranial ectoderm towards ear identity.

## Introduction

In the developing embryo, cellular diversity arises through a series of cell fate decisions. Understanding how these decisions take place is therefore a central objective of developmental biology. Cell fate choice is mediated by regulatory factors, which activate a set of transcription factors that in turn control the expression of proteins required for cell-specific functions. Direct lineage reprogramming has emerged as a ground-breaking concept, allowing cells to switch fates whilst bypassing pluripotency, a ‘shortcut’ aimed at improving the speed and efficiency of cell fate conversion (1-3). In turn, lineage reprogramming also highlights the central role of regulatory factors in determining cell fate. Classical examples are the transcription factors MyoD, which can reprogram fibroblasts into myogenic cells (4), and Pax6 which induces ectopic eyes when mis-expressed in non-eye cells (5-7). Fundamental for the perception and interaction with their environment, sense organs and their diversification have enabled vertebrates to thrive in almost every environmental niche. However, apart from Pax6 in the eye, key regulators for other sense organs have not yet been discovered.

Like the eye, the inner ear is a pan-vertebrate sense organ and is responsible for the perception of sound and movement (8-11). During development, it arises from a shared pool of progenitors which also gives rise to epibranchial neurons. These otic-epibranchial progenitors (OEPs) reside next to the cranial neural plate where they are intermingled with neural and neural crest precursors (12, 13). Subsequent signaling from adjacent tissues induces the segregation of these fates into distinct territories (14-21). While epibranchial cells produce sensory ganglia, the otic placode invaginates to form a vesicle, which is then transformed into the inner ear, containing many specialized cell types and associated neurons. While signals conferring inner ear identity have been extensively studied, we still lack a comprehensive understanding of the epigenetic mechanisms and transcription factors regulating its specification.

Here, we model gene expression dynamics during the segregation of otic and epibranchial fates to identify key regulators of ear fate. We use single-cell-RNA-sequencing (scRNAseq) to ask whether OEPs are progenitors with mixed identity or pre-biased towards their later fate and pseudo-time analysis to model transcriptional changes during ear specification. Using epigenomic profiling we provide the first genome-wide identification of ear enhancers and their upstream regulators. Together with functional experiments, we identify a small transcriptional circuit that defines ear identity comprising Sox8, Pax2 and Lmx1a, with Sox8 at the top of the hierarchy. Sox8 alone triggers the ear program in ectodermal cells and initiates ear morphogenesis by forming ear vesicles containing differentiating neurons. Thus, using a multi-omics approach we have uncovered Sox8 as a critical ear fate determinant and potential reprogramming factor within the developing cranial ectoderm.

## Results

### Dynamic changes in gene expression characterize the transition from progenitor to ear commitment

To unravel the genetic hierarchy that controls how otic and epibranchial cells diverge, we first characterized the transcriptional profile of cells committed to each fate (22) (Fig.1 A-D; Fig. S1A-F). To label each cell population we used two enhancers driving EGFP, the novel otic Lmx1aE1 enhancer (see below) and the epibranchial enhancer Sox3U3 (23). Reporter constructs were electroporated into the OEP territory of somite stage (ss) 3-5 embryos. At ss18-21, the ear region was dissected and EGFP^+^ cells were isolated by fluorescence-activated cell sorting (FACS) and processed for RNA-seq. Differential expression analysis identifies 103 and 319 genes upregulated in Lmx1aE1-EGFP and Sox3U3-EGFP expressing cells, respectively (log2FC > 1.5; adjusted p-value < 0.05; Fig. 1D; Fig. S1C, D and Supplementary Data 1). This analysis defines the transcriptional states of definitive otic and epibranchial cells.

**Figure 1.**
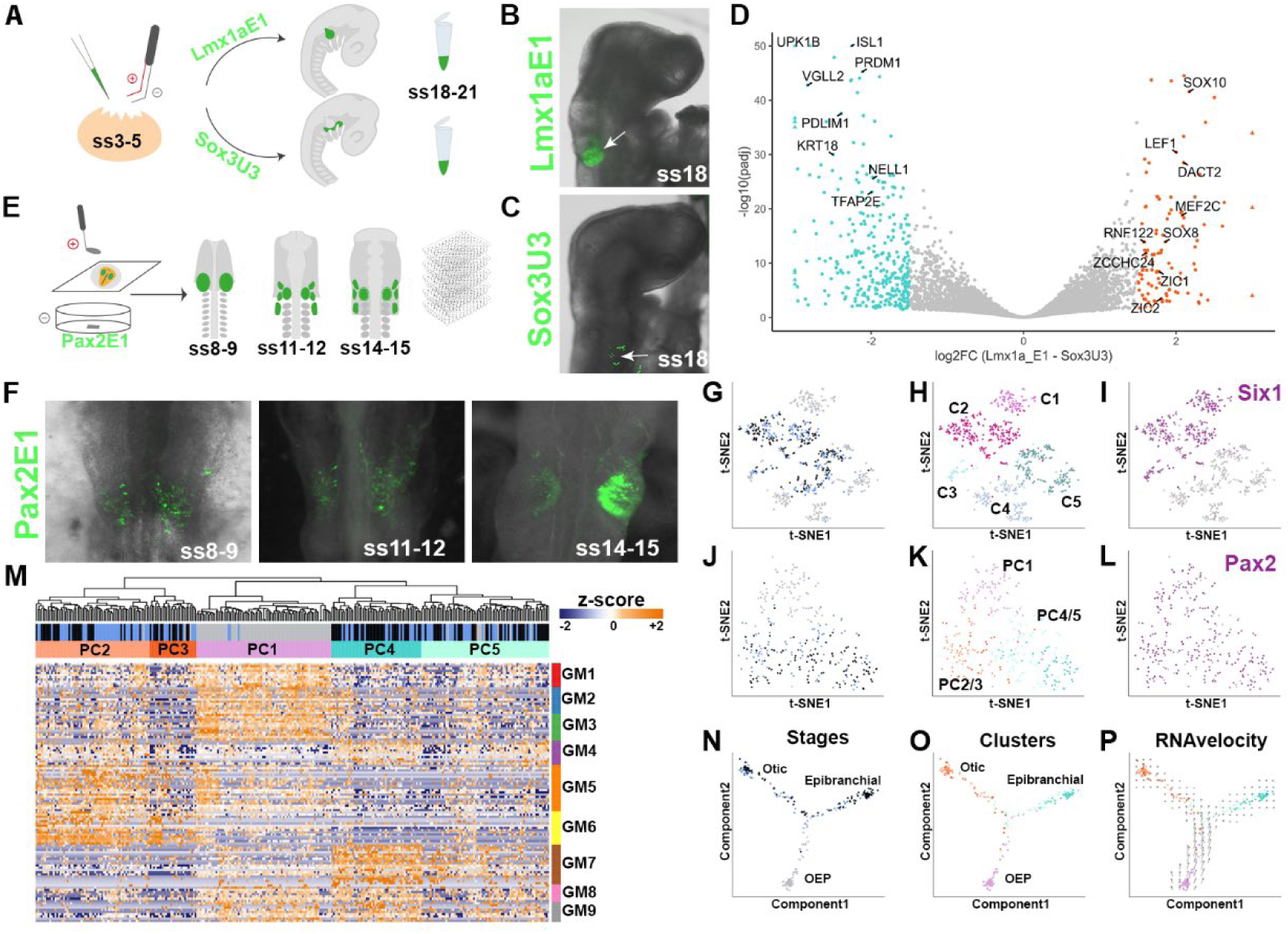
Transcriptomic characterization of ear development. (**A-C**) *In ovo* electroporation (**A**) was used to label and collect otic (**B**; Lmx1aE1-EGFP^+;^ white arrow) and epibranchial (**C**; Sox3U3-EGFP^+^; white arrow) cells for bulk RNAseq. (**D**) Volcano plot showing genes differentially expressed (absolute log2 fold-change >1.5 and adjusted p-value < 0.05) in otic (orange) and epibranchial cells (green). (**E-F**) Cells expressing the Pax2E1-EGFP reporter active in OEPs, otic and epibranchial placodes were collected for scRNAseq at the stages indicated (**F**). (**G**-**L**), tSNE representation of unsupervised hierarchical clustering of all cells (**G-I**) and of the placodal cell subset (**J-L**); cells were color-coded according to the stage collected (**G, J**; OEP grey, ss11-12 blue, ss14-15 black), clusters (**H, K**) and placodal marker expression: *Six1* (**I**) and *Pax2* (**L**). (**M**) Heatmap showing partitioning of the placodal subset (C1, C2 in **H**) into five clusters (PC1-5) based on gene modules (GM). Expression profiles reveal that PC1 largely contains OEP-like cells, while PC2/3 and PC4/5 are composed of otic-like and epibranchial-like cells. (**N**-**O**), Pseudo-time ordering using Monocle2 shows trajectories between stages (**N**) and clusters (**O**). Note: otic- and epibranchial-like cells segregate into two branches: otic-like in orange, epibranchial-like in green, OEP-like in pink. (**P**), RNAvelocity vector field verifies the directional trajectories predicted by Monocle2.

Next, we investigated the transcriptional changes that take place as OEPs transition from a common progenitor population to definitive otic and epibranchial cells. *Pax2* is expressed throughout this time-window in both cell populations (Fig. S1G, H) (12, 22). Using a Pax2E1-EGFP reporter (see below), we isolated single cells by FACS from consecutive stages of ear specification: OEP (ss8-9), early-placode (ss11-12) and late-placode stage (ss14-15) and processed them for scRNAseq (Fig. 1E, F; Fig. S1I, S2B-D). To characterize cellular diversity, we first looked at groups of genes co-expressed across the dataset (gene modules). These gene modules were identified in an unbiased manner through hierarchical clustering of a gene-gene Spearman correlation matrix. Gene modules of interest were selected based on the presence of well-characterized makers for placodal, neural and neural-crest cells including the new otic and epibranchial genes identified (Fig. 1D). Initial cell clustering defines five major clusters (Fig. S2A) to which we assigned identities using known markers (Fig. 1G-H, Fig. S2F-I). Clusters C1 and C2 represent *Pax2*^*+*^*/Six1*^*+*^ cells expressing high levels of OEP and placodal makers (Fig. 1H, I; Fig. S2F, I), while cluster C3 contains contaminating mesoderm (*Twist1*^*+*^, *Sim1*^*+*^; Fig. S2I). Surprisingly, we also find two clusters with low levels of *EGPF* mRNA (Fig. S2A, E) and relatively few *Pax2*^*+*^ cells: one containing neural-like cells (C4; Sox21^+^) and another containing neural crest-like cells (C5; Pax7^+^; Fig. S2A, G, H, I). Since OEPs are mixed with future neural and neural crest cells at ss8-9 in a *Pax2+* territory (12, 24) and these precursors can co-express markers for different fates prior to differentiation (25), this observation suggests that while cells in clusters C4/5 initially activate Pax2E1-EGFP they subsequently downregulate enhancer activity and *Pax2* expression.

To investigate the transcriptional dynamics accompanying otic and epibranchial fate decisions, we subset the placodal clusters (C1/2 in Fig. 1; Fig. S2A). Re-clustering these cells using gene modules containing otic and epibranchial genes, we obtained five placodal clusters (PC1-5; Fig. 1M, J, K; Supplementary Data 2). PC1 largely consists of only OEPs (cells collected at ss8-9), while the other clusters contain cells from both early and late placode stages (Fig. 1M, J, K). Indeed, PC1 is characterized by the expression of OEP genes (GM1-3; Fig. 1M), as well as sharing genes with the otic module GM5/6 and the epibranchial module GM7-9. In contrast, PC2/3 and PC4/5 are transcriptionally distinct from each other with profiles akin to otic (GM5/6) and epibranchial (GM7-9) cells, respectively. To explore the relationship between different cell clusters we organized cells along pseudo-time using Monocle2 (26). This analysis predicts that OEPs gradually split into one otic and one epibranchial branch, each composed of early and late placodal cells (Fig. 1N, O). Calculating RNA-velocity independently (27) and embedding the corresponding vector field onto the Monocle2 trajectory validates the directionality of predicted cell state transitions (Fig. 1P).

To explore dynamic changes of gene expression accompanying these inferred trajectories, we used Branch-Expression-Analysis-Modelling (BEAM) (26). This identifies groups of transcription factors expressed in OEPs prior to the branching point, which subsequently segregate into either the otic (e.g. *Sox8, Lmx1a, Pax2, Zbtb16*) or the epibranchial (e.g. *Foxi3 (28), Tfap2a/e, Nell1*) branch (Fig. 2A, Fig. S3A). To quantify the changes in co-expression of otic and epibranchial genes, we assessed the proportion of co-expressing cells before and after the branching point. A two-tailed Wilcoxon rank sum test reveals significantly more cells co-expressing otic and epibranchial markers in OEPs than in epibranchial (W=214, p=0.0013) and otic cells (W=235, p<0.0001) after the branching point (Fig. 2B, C). Quantification of gene expression in the monocle trajectories and by *in situ* hybridization chain reaction (HCR) (29) confirms that otic (*Sox8, Lmx1a*) and epibranchial (*Foxi3, Tfap2e*) transcripts overlap at ss8-9, and that their expression resolves as both placodes are firmly established (Fig. 2D-I, Fig. S3B-H).

**Figure 2.**
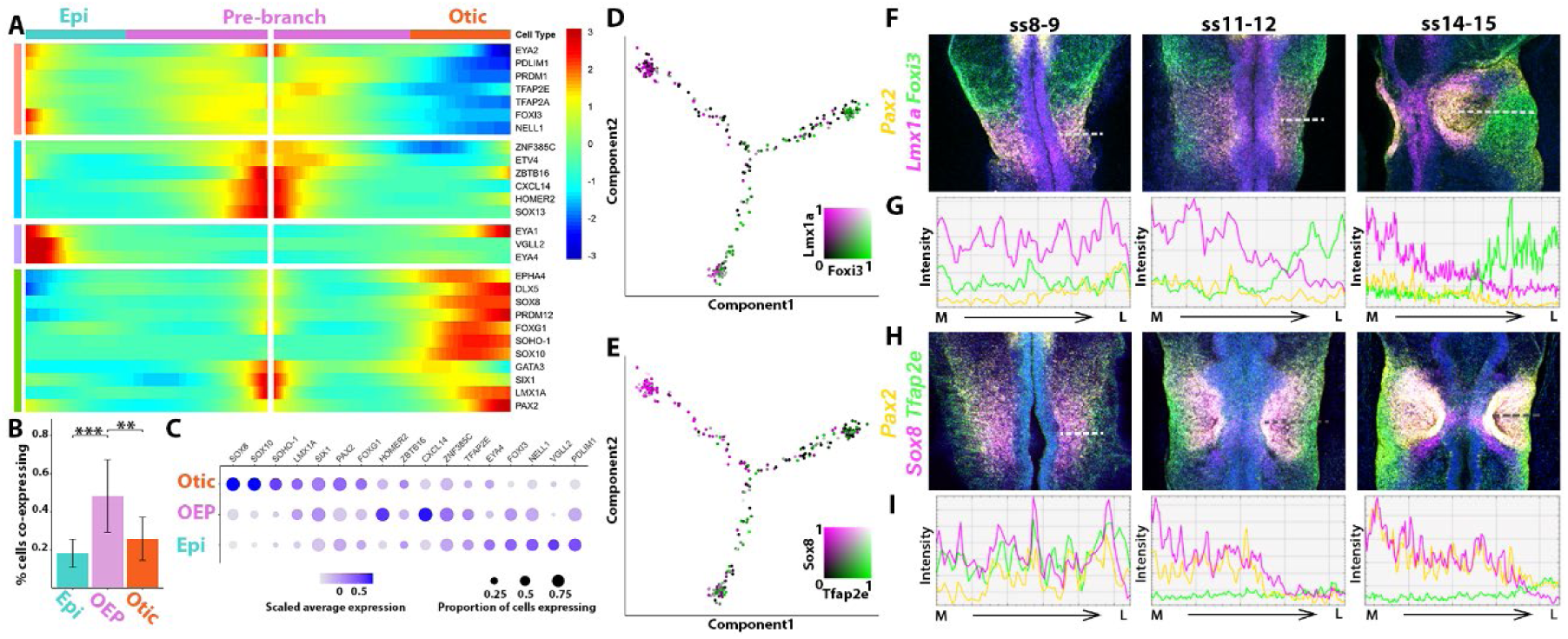
Dynamic gene expression as OEPs segregate into otic and epibranchial fates. (**A**) BEAM identifies genes regulated in a branch-dependent manner. (**B**) Histogram showing the proportion of cells co-expressing genes that are expressed before the branching point, but later segregate to the otic or epibranchial branch. Significantly more cells co-express such genes before the branching point than thereafter. Error bars indicate +/- 1 standard deviation. ** p-value < 0.01; *** p-value < 0.001. (**C**), Dot plot for OEP, otic and epibranchial markers based on scRNAseq data. Expression level is indicated by color intensity and gene expression frequency by dot size. (**D**-**E**), A proportion of cells co-expresses otic (*Sox8, Lmx1a*) and epibranchial (*Foxi3, Ap2e*) markers prior to the branching point. (**F**-**I**) *In situ* HCR (**F, H**) and intensity measurements (**G, I**) along the medial (M) to lateral (L) axis of the OEP-domain indicated by a dashed line in **F, H**, confirms gradual segregation of OEP gene expression as distinct otic and epibranchial cell populations emerge.

Together, these results identify groups of transcription factors whose expression changes over time as otic and epibranchial precursors segregate and therefore may play a key role in cell fate decisions.

### Epigenomic profiling uncovers new regulatory elements and motifs in ear precursors

Transcription factors controlling cell fate choice regulate their downstream targets by interacting with tissue specific cis-regulatory enhancer elements. Active enhancers are regions of open chromatin flanked by nucleosomes enriched for histone 3 lysine-27 acetylation (H3K27ac), while actively transcribed genes are marked by H3K4me3 (30-33). We therefore profiled ss8-9 OEPs by ChIPseq for H3K27ac, H3K4me3 and the repressive mark H3K27me3 and determined chromatin accessibility by ATACseq. Overlapping H3K27ac and ATACseq data identifies 10969 genomic regions that also show depleted H3K27me3 marks; average profiles show bimodal H3K27ac read distribution surrounding ATACseq peaks (Fig. S4A, B). Of these just over 70% are intergenic or intronic representing putative enhancers (8316), while the remaining are close to transcription start sites (TSS; Fig. S4C). We associated each putative enhancer to the nearest TSS of protein coding genes; GO term analysis of the corresponding genes returns MAP-kinase, Wnt and Notch signaling known to mediate ear induction, development and neurogenesis (Fig. S4D) (16, 34). To assess their activity in vivo we selected putative enhancers in the vicinity of ear-enriched genes, generated EGFP reporters and co-electroporated them with ubiquitously expressed mCherry into head-fold-stage chick embryos. RT-PCR-based assays (35) (not shown) and fluorescence microscopy confirm enhancer activity in ear progenitors and otic placodes (Fig. 3A, B; Fig. S5-7). To identify upstream regulators that may act as otic determinants, we performed motif enrichment analysis of all 8316 putative ear enhancers. This reveals an over-representation of binding sites for Sox, TEAD and Six family members and for Tfap2a (Fig. 3C; Fig. S4E). Of these, Tfap2a and Six proteins have previously been implicated in cranial placode development (36-40), confirming that this strategy can identify relevant regulatory factors.

**Figure 3.**
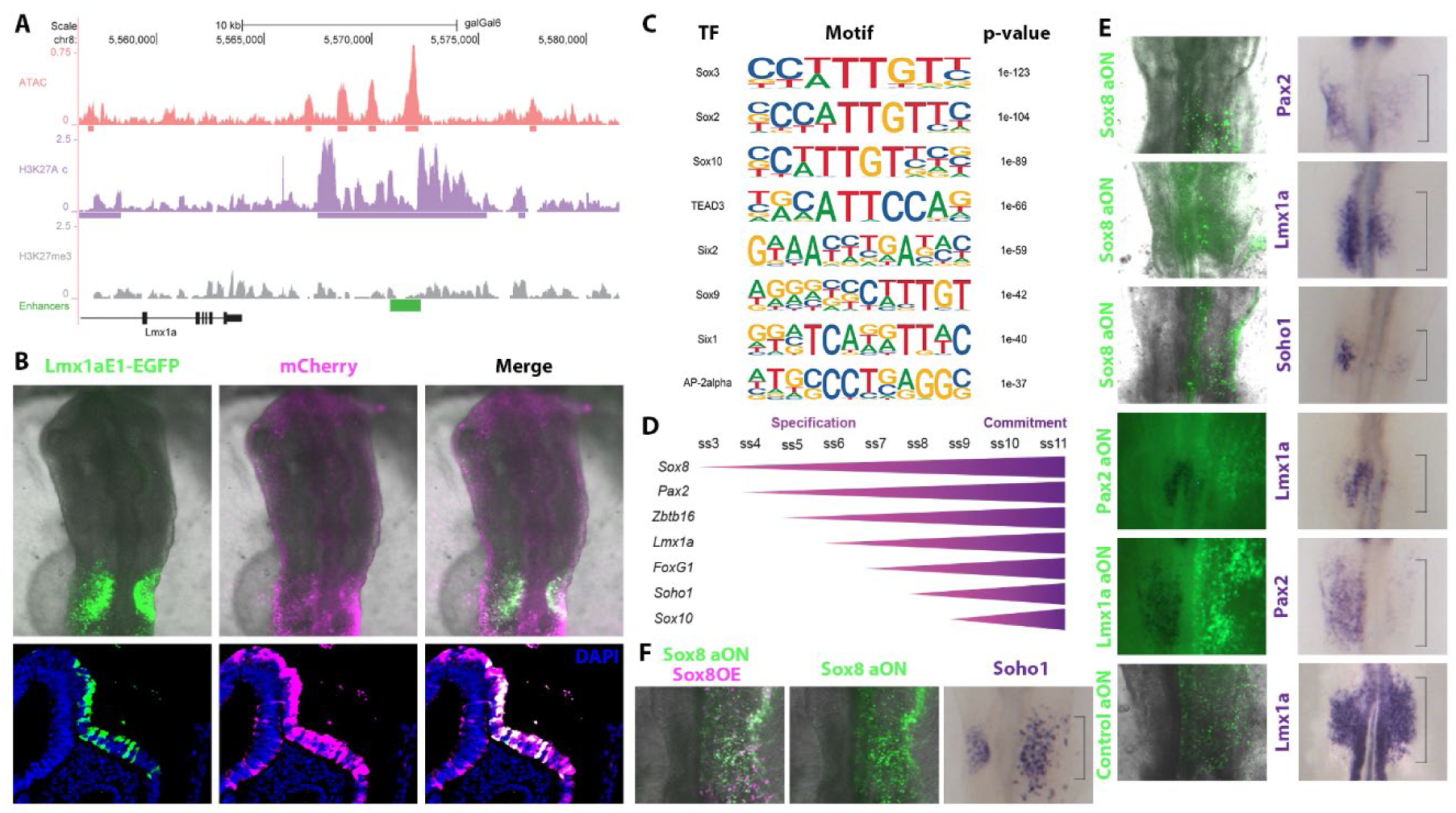
Identification of putative regulatory regions in OEPs. (**A**) Genome browser view of ATAC, H3K27ac and H3K27me3 profiles in OEPs at the Lmx1a locus. Cloned putative enhancer is shown in green. (**B**) Co-electroporation of Lmx1aE1-EGFP reporter (green) and constitutive mCherry (magenta) reveals *in vivo* enhancer activity in the otic placode. Sections reveal that the enhancer activity is restricted to the ectoderm at the level of the otic placode: Lmx1aE1 in green, constitutive mCherry in magenta and DAPI in blue. (**C**) Motif enrichment analysis of all identified enhancers. (**D**) Diagram showing the onset of expression of potential otic regulators during specification and commitment stages. (**E**) Unilateral knock-down of selected transcription factors using fluorescein labelled antisense oligonucleotides (aON; green) leads to downregulation of otic markers as shown by *in situ* hybridization (purple) on the targeted side of the embryo. Note: fluorescent images are taken prior to *in situ* hybridization for Sox8aON and controls. (**F**) Unilateral co-electroporation of Sox8aON (green) and Sox8-mCherry construct (magenta) on the right side of the embryo restores *Soho1* expression in the otic territory. Brackets on the right indicate the normal size of the otic region.

We also exploited the idea that cell identity genes may be regulated by super-enhancers characterized by high density of H3K27ac, while their gene bodies are decorated with H3K4me3 (41-43). Examining OEP transcription factors that segregate to the ear lineage (Fig. 2A), we find that the Sox8 locus is marked with broad H3K4me3 (Fig. S4F), while enhancers close to Lmx1a, Zbtb16 and Sox13 are putative super-enhancers (Fig. 3A, Figs. S4F, S6A, S7A). These results identify new regulatory elements that control gene expression during early ear development as well as several factors that may act as otic specifiers.

### Defining core components of the ear determination network

Together, the BEAM and epigenomic analysis point to Lmx1a, Zbtb16 and members of the Sox family as potential regulators of ear identity, while previous studies have also implicated Pax2 (44). We next examined the temporal sequence of their expression in otic progenitors. Of the Sox genes, *Sox3* and *Sox13* are expressed prior to ear specification (45, 46), while *Sox9* and -*10* are activated later (45, 47). These genes are therefore unlikely to initiate the otic program. In contrast, the SoxE group factor *Sox8* is highly enriched in OEPs prior to segregating to otic cells (Fig. 3D). *In situ* hybridization reveals that *Sox8* begins to be expressed at 3ss, followed shortly thereafter by *Pax2, Zbtb16* and *Lmx1a*, while the known otic factors *Foxg1, Soho1* and *Sox10* are activated later (Fig. 3D, Fig. S8A).

To explore the regulatory interactions between the earliest OEP transcription factors, we systematically knocked down each one and assayed the expression of all others as well as that of *Foxg1* and *Soho1* as a readout for otic identity using *in situ* hybridization and RT-qPCR (Fig. 3E; Fig. S8B, C, Fig. S9A). Control or antisense oligonucleotides targeting Sox8, Pax2, Zbtb16 or Lmx1a were electroporated into future OEPs of head-fold-stage chick embryos and gene expression was analyzed at OEP stages (ss8-9). We find that Sox8 is necessary for the expression of all assayed ear transcription factors. Pax2 is required for the expression of *Zbtb16* and *Lmx1a*, which in turn are necessary for *Pax2*, suggesting that they act in a positive feedback loop with Pax2. Zbtb16 is also necessary for *Foxg1*. All gene expression changes can be rescued by co-electroporation of the appropriate full-length constructs (Fig. 3F, Fig. S9B-E). Furthermore, transcription factor binding site analysis identifies motifs for the core ear-network factors (Sox8, Pax2, Lmx1a, Zbtb16) in enhancers associated to *Lmx1a, Pax2* and *Zbtb16* (Fig. S4G, Supplementary Data 3, 4) suggesting that these interactions might be direct. In summary, *Sox8* is the earliest OEP transcription factor and later becomes confined to otic cells. Our functional experiments put Sox8 at the top of the ear determination network forming a regulatory circuit with Pax2, Lmx1a and Zbtb16.

### Sox8 induces ectopic otic vesicles and vesicle-derived neurons

If these factors indeed form a minimal circuit driving otic specification, they should be able to convert non-ear cells into cells with ear identity. We tested this hypothesis by electroporating different combinations of Sox8-, Pax2-, Lmx1a- and Zbtb16-mCherry-tagged constructs into head-fold-stage ectoderm not destined to contribute to the ear together with the Lmx1aE1-EGFP reporter. We find that misexpression of all four factors and of Sox8/Pax2/Lmx1a activates robust expression of the reporter, while combinations lacking Sox8 do not (Supplementary Table7). In addition, Sox8/Pax2/Lmx1a electroporation also results in the formation of many *Soho1*^*+*^ otic vesicles scattered across the head ectoderm as well as neurofilament positive neurons (Fig. 4A), the first cell type to differentiate in the otic vesicle (48, 49).

**Figure 4.**
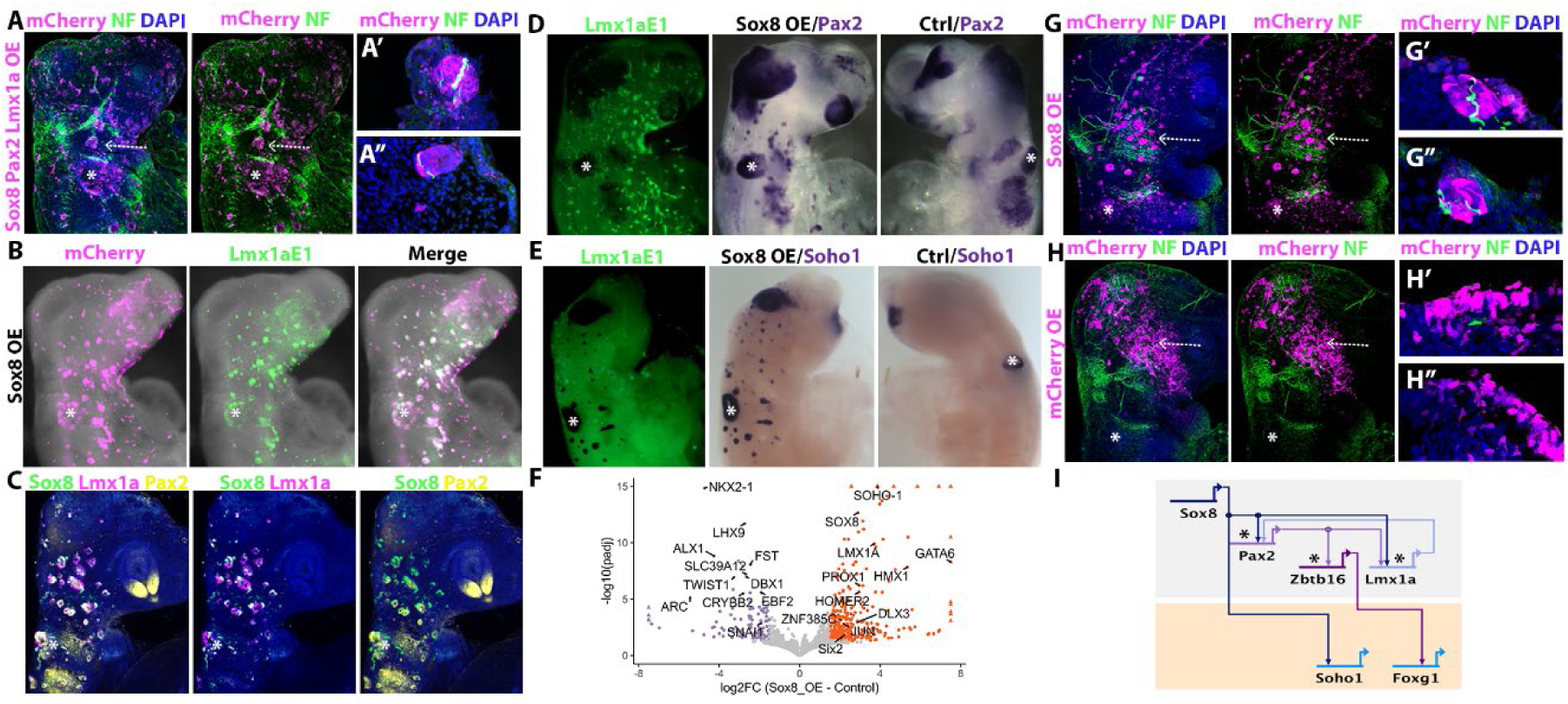
Sox8 converts non-otic cells into otic vesicles with associated neurons. (**A**) Co-electroporation of mCherry-tagged vectors containing the full-length sequence of Sox8/Pax2/Lmx1a (magenta) generates ectopic vesicles across the cranial ectoderm associated with neurofilament positive neurons (green). (**A’, A’’**) Transverse sections at the level indicated by arrows in **A** show neurofilament/mCherry positive neuronal projections from the ectopic vesicles (white). (**B**) Overexpression of Sox8 (Sox8 OE; magenta) alone activates Lmx1aE1-EGFP (green). (**C**-**E**) Overexpression of Sox8 (Sox8 OE) induces otic markers on the electroporated, but not on the control side (Ctrl) as shown by HCR (**C**) and *in situ* hybridization (**D, E**) for otic markers. (**F**) RNAseq of Sox8 and control electroporated cranial ectoderm; volcano plot shows enrichment of otic genes after Sox8OE (absolute log2 fold-change >1.5 and adjusted p-value < 0.05). (**G**) Sox8-mCherry overexpression (magenta) generates ectopic otic vesicles with neuronal projections (green), while controls do not (**H**); transverse sections of ectopic otic vesicles (**G’, G’’**) and control ectoderm (**H’, H’’**) show neurofilament positive neurons within the ectopic vesicle. White asterisks (**A**-**H**) indicate the endogenous otic vesicle. (**I**) BioTapestry model showing the minimal transcriptional circuit for otic specification with Sox8 at the top of the hierarchy. Black asterisks indicate enhancers with predicted Sox8 binding motifs.

We next asked whether Sox8 alone can initiate the ear program. We find that misexpression of Sox8-mCherry alone activates the Lmx1aE1-EGFP reporter in ectopic vesicles (Fig. 4B), as well as the expression of *Pax2, Lmx1a* and *Soho-1* (Fig. 4C-E; Fig. S10A). To assess to what extent Sox8 can confer ear identity, we isolated double-positive Sox8-mCherry/Lmx1aE1-EGFP cells from ss11-12 by FACS and compared their transcriptome with control ectoderm labelled with constitutive mCherry/EGFP. Differential expression analysis shows upregulation of 399 transcripts in comparison to controls, while 112 genes are downregulated (log2FC > 1.5; adjusted p-value < 0.05) (Fig. 4F, Fig. S10B-C, Supplementary Data 5). Seventeen of the twenty seven upregulated transcription factors are known to be expressed in the inner ear, while for the remaining ten no expression data are available (Supplementary Data 6). In contrast, among the downregulated genes are typical neural crest and forebrain transcripts. To confirm that Sox8 expressing cells have indeed acquired otic character we assessed the expression of otic enriched TFs from previously published data sets (45) in Sox8 overexpressing and control cells (Fig. S10E). Of 110 otic TFs, 98 are upregulated after Sox8 misexpression, but not in controls. This observation suggests that Sox8 alone can confer ear identity to cranial ectoderm cells. Can Sox8 alone also trigger neurogenesis? Indeed, Sox8-induced ectopic vesicles are associated with neurofilament-positive neurites generated from Sox8 expressing cells themselves, while control cells do not form vesicles or neurons (Fig. 4G, H).

Together, our results position Sox8 at the top of the otic GRN (Fig. 4I), modulating the activity of other ear factors like *Pax2, Lmx1a* and *Soho1*. Downstream of Sox8, Lmx1a and Pax2 seem to form a positive feedback loop with Pax2 required for *Zbtb16* activity which in turn regulates *FoxG1*. Thus, Sox8 can activate the transcriptional program for ear fate in cells destined to form other sense organs or epidermis.

## Discussion

In this study, we have identified critical components of the ear determination network: Sox8, Pax2 and Lmx1a (Fig. 4I). To do so we have used different criteria all of which converge on these three transcription factors: i) temporal sequence of expression; ii) segregation of expression to otic fate using pseudo-temporal ordering of single cells and BEAM; iii) position at the top of the otic gene network (45); iv) motif enrichment of newly identified enhancers; v) association with histone marks that define super-enhancers and/or fate determinants and finally vi) requirement for the expression of known ear markers.

Our analysis puts Sox8 at the top of the transcriptional hierarchy that controls ear fate; Sox8 alone imparts ear identity to cells otherwise destined to form head epidermis, other sense organs or cranial ganglia. Previous findings have implicated Spalt4 and other SoxE group family members in otic vesicle formation. However, their misexpression generates only few ectopic vesicles next to the endogenous ear. Unlike Sox8, Spalt4 and Sox10 cannot activate neurogenesis and neuronal differentiation (50, 51) pointing towards a limited ability of these factors to reprogram ectoderm into functional ear cells. Together, these findings highlight the importance of SoxE group TFs during inner ear development and propose a prominent role for Sox8 in activating the transcriptional program for the inner ear. Analysis of ear-specific regulatory elements (Supplementary Table 3) suggests that many downstream effectors are directly activated by Sox8. In future, Sox8 may serve as a key factor for rapid cell conversion in the context of regeneration and repair in the inner ear.

While Sox8 regulates ear-specific enhancers (52-54) (this study), Pax2 is involved in different steps of otic placode formation and patterning (44, 55-57). Sox proteins cooperate with a variety of transcription factors to exert their cell type specific function (58). It is therefore tempting to speculate that, in analogy to the eye, where Sox2 cooperates with Pax6 to regulate lens-specific transcription (59), in the ear Sox8 might partner with Pax2.

Finally, our results also show that OEPs initially express competing transcriptional programs that resolve over time as otic and epibranchial cell states are established. We capture previously unknown gene modules that accompany this process as well as regulatory regions associated with otic-epibranchial specification. In turn, this information is critical to unravel the underlying gene regulatory networks and identify the transcription factor codes that determine cell identity in the cranial sensory nervous system. In the long term, this will enhance our ability to engineer specific sensory cell types for basic research and regenerative purposes.

## Materials and Methods

### Expression and enhancer constructs

Putative enhancers were amplified from chick genomic DNA and cloned into pTK-EGFP reporter vectors after digestion with XcmI (35). To generate expression constructs total RNA was isolated from HH8-12 chick embryos with RNAqueous-Micro Total RNA Isolation Kit (Thermo Fisher Scientific), reverse transcribed using Superscript II reverse transcriptase (Thermo Fisher Scientific, 180644-014) and oligo-dT primer. Specific primers were used to amplify the full-length coding sequence of Sox8, Pax2, Lmx1a and Zbtb16 and PCR products were cloned into pCAB-IRES-mCherry. All sequences were verified by Sanger sequencing.

### Chick embryos, electroporation and culture

Fertilized hens’ eggs (Stewart, Norfolk UK) were incubated at 38°C and staged according to Hamburger and Hamilton (HH) (60). All experiments were performed on embryos younger than 12 days, thus were not regulated by the Animals Scientific Procedures Act 1986.

OEPs for bulk RNAseq were labelled using *in ovo* electroporation (61); eggs were incubated until the 3-6 somite stage (ss), pTK-Lmx1aE1-EGFP or pTK-Sox3U3-EGFP plasmids (1μg/μl) were injected targeting the OEP territory. Electroporation using Ovodyne electroporator (TSS20, Intracel) was performed with five 50ms pulses of 8V at 100ms intervals. After incubation until ss18-21, embryos were collected in PBS and processed for bulk RNAseq.

For *ex ovo* culture, embryos were harvested using filter paper rings (62) and cultured on egg albumen; for long-term culture (48hrs) the ‘modified Cornish pasty’ method (63) was used. *Ex ovo* electroporation was performed to collect cells for scRNAseq, for knock-down, rescue and overexpression experiments. The posterior placodal region or the cranial surface ectoderm of HH6 embryos was targeted for electroporation by injecting plasmid DNA at 1μg/ul (pTK-Pax2E1-EGFP to label OEPs; pCES-Sox8-mCherry + pTK-Lmx1aE1-EGFP, pCES-Sox8-mCherry + pCES-Lmx1a-mCherry + pCES-Pax2-mCherry; pCES-mCherry + pCES-EGFP for misexpression), control or antisense-oligonucleotides (aON; GeneTools; 0.75mM with 0.3μg/μl carrier DNA), or a combination of aON and expression construct (0.75mM aON + 0.75μg/μl plasmid). For knock-down and rescue experiments, unilateral injections on the right side of the embryo were performed while the uninjected contralateral left side served as an internal control. Oligonucleotides used were Sox8: 5’-CTCCTCGGTCATGTTGAGCATTTGG-3’(51), Pax2: 5’-GGTCTGCCTTGCAGTGCATATCCAT-3’(49), Lmx1a: 5’-CCTCCATCTTCAAGCCGTCCAGCAT-3’(42), Zbtb16: 5’-GTCAAATCCATAGCACTCCCGAGGT-3’, Sox13: 5’-CTCCTCATGGACATCCATTTCATTC-3’ and control 5′-ATGGCCTCGGAGCTGGAGAGCCTCA-3′. Electroporation was performed in a chamber using five 5V pulses of 50ms in 100ms intervals (57).

To monitor fluorescence, electroporated embryos were imaged using a Zeiss Axiovert 200M microscope with a Hamamatsu C4742-95 camera and using HC image software. Fluorescent images were taken prior to processing for *in situ* hybridization and antibody staining.

### Whole mount and hybridization chain reaction fluorescent in situ hybridization

*In situ* hybridization was carried out following previously described protocols (64). Whole-mount pictures were taken using an Olympus SZX12 with a Retiga2000R camera and Q-Capture Pro7 software. Paraffin embedded embryos were sectioned at 8μm sections in a Leica RM2245 microtome. Upon sectioning, images were taken in a Zeiss ApoTome.2 coupled with an Axiocam 503 color camera and using the ZEN 2.5 software.

HCR v3 was performed using the Molecular Technologies protocol (29). Briefly, embryos were fixed in 4% PFA for one hour at room temperature, dehydrated in a series of methanol in PBT and stored overnight at -20°C. After rehydration and proteinase-K treatment (20mg/ml; 3 min) embryos were post-fixed in 4% PFA for 20 min. Embryos were then washed on ice in PBS, 1:1 PBT/5X SSC (5X sodium chloride sodium citrate, 0.1% Tween-20) and 5X SSC for 5 min each. Pre-hybridization in hybridization buffer was performed for 5 min on ice, followed by 30 minutes at 37°C. Embryos were hybridized overnight at 37°C with probes at 4pmol/ml in hybridization buffer. After four 15 min washes with probe wash buffer at 37°C, preamplification was carried out in amplification buffer for 5 min at room temperature. Hairpins were prepared individually at 30pmol final concentration; they were incubated at 95°C for 90 seconds followed by cooling to room temperature for 30 min, protected from light. Cooled hairpins were added to 500μl amplification buffer, embryos were incubated in hairpins overnight at room temperature followed by two 5 min and two 30 min washes in 5X SSC. After a 5 min incubation in DAPI (10 mg/ml), they were washed three times for 10 min with 5X SSC, before being imaged using Leica SP5 laser scanning confocal inverted microscope using the LAS AF software.

For HCR image analysis, Z-stacks were collected for 50-70μm; figures show projections of all stacks. Images were processed using ImageJ and the ImageJ Plot Profile tool was used to calculate intensity plots. In brief, an optical section in the centre of the placode territory was selected using the Pax2 channel as a reference. The Pax2 channel was added to the Sox8, Lmx1a, Foxi3 or Tfap2e channels, respectively. Intensity values were then calculated across the area of interest and plotted.

### Whole mount immunostaining

Embryos were collected in PBS, fixed for 25 minutes at room temperature in 4% PFA, washed in PBS-Tx (PBS + 0.2% Triton X-100) and blocked in 5% goat serum in PBS-Tx for 3–5 h at room temperature. Embryos were then incubated in primary antibody diluted in blocking buffer for 24-72 hours at 4°C. Primary antibodies were rabbit anti–mCherry (1:200; Abcam ab167453), mouse anti– NF (1:100; Thermo Fisher Scientific 13-0700), rabbit anti-GFP (1:500; Abcam a11122) or mouse anti GFP (1:1000; Molecular Probes A11120). After five 60 min washes and one overnight wash in PBS-Tx, embryos were incubated in secondary antibodies (1:800) at 4°C overnight. Secondary antibodies used were goat anti–rabbit IgG Alexa Fluor 488 (Thermo Fisher Scientific, A11036), donkey anti–rabbit Alexa Fluor 568 (Thermo Fisher Scientific, A11001), goat anti mouse IgG Alexa Fluor 488 (Molecular Probes A11001) and goat anti mouse IgG Alexa Fluor 568 (Molecular Probes A11004). Embryos were then briefly incubated in PBS containing 10 mg/ml DAPI and washed at least five times in PBS-Tx before being mounted on slides and imaged using Leica SP5 laser scanning confocal inverted microscope using a 10x objective or an Olympus SZX12 with a Retiga2000R camera and Q-Capture Pro7 software. Confocal whole mount images in the manuscript are maximum intensity projections of embryo z-stacks. Sections were imaged using a 63x oil immersion objective and maximum intensity projections are shown.

### Cryosectioning

Embryos were embedded in gelatine as previously described (65) and cryo-sectioned at 15-20μm using a Bright OTF5000 cryostat. Sections were mounted using Mowiol 4-88 (Sigma Aldrich, 81381) and imaged using Leica SP5 laser scanning confocal inverted microscope (LAS AF software) or a Zeiss Axiovert 200M microscope with a Hamamatsu C4742-95 camera and using OCULAR software.

### FAC-Sorting of cells

Cells were collected using a BD FACS-Aria Fusion. For single-cell-RNA-seq three batches of experiments were performed, one per stage. Live EGFP+ cells were selected using propidium iodide (for ss8-9 cells) or DAPI (ss11-12, ss14-15) as a live/death cell marker. The gating tree was set as follows: first FSC-A/SSC-A which represents the distribution of cells based on size and intracellular composition, respectively. Then either FSC-A/FSC-H (ss8-9) or SSC-W/SSC-A (ss11-12, ss14-15) was used to exclude the events that might represent more than one cell. Next, we performed a live gate to select the cells that were propidium iodide/DAPI negative. Finally, GFP+ cells were identified and selected for sorting. For bulk experiments we used DAPI as a live/death marker and gating was performed as described above for ss11-12/ss14-15. For Sox8OE/Lmx1a-E1+ and mCherry/EGFP+ control cells the last step of the gating tree was performed using GFP/mCherry to select the double positive population. In all the experiments a 100-micron nozzle and 20psi pressure was used.

### Quantitative PCR (qPCR)

RNA from dissected otic tissue was isolated using the RNAqueous-Micro Kit (Ambion, AM1931) and reverse transcribed. Primers for target genes were designed with PrimerQuest (IDT). qPCR was performed using Aria Mx Real-Time System (Agilent Technologies) with SYBR green master mix (Roche, 64913850001). RT-qPCR for antisense oligonucleotide experiments was carried out with a minimum of three biological replicates; two sample Welch’s t-tests with Benjamini-Hochberg multiple test correction were used to determine statistical significance between control and antisense oligonucleotide ΔCt values. Relative expression (2-ΔΔCt) (66) was calculated using the NormqPCR package in R. The geometric mean of Gapdh and Rplp1 expression, or Rplp1 expression alone, was used to normalize gene expression.

### Bulk RNA sequencing

To label otic and epibranchial cells embryos were electroporated with Lmx1aE1-EGFP and Sox3U3-EGFP plasmids, respectively, and whole heads were used for cell collection. For overexpression experiments embryos were electroporated with Sox8-mCherry+Lmx1aE1-EGFP or with pCAB-mCherry+pCAB-EGFP, the endogenous otic placode and the trunk were removed before cell dissociation. Cells were dissociated in FACSmax cell dissociation solution (Ambion, T200100) containing papain (30U/mg, Sigma-Aldrich, 10108014001) for 20min at 37°C before being transferred to Hanks Balanced Solution without calcium and magnesium (HBSS, Life Technologies, 14185045) containing 5% heat-inactivated foetal-bovine-serum (FBS), rock inhibitor (10 μM, Stemcell Technologies, Y-27632) and non-essential amino acids (Thermo Fisher Scientific, 11140035). Cells were disaggregated by pipetting, sequentially filtered through 0.35μm and 0.20μm filters (Miltenyi Biotech, 130-101-812). Pelleted cells were resuspended in 500μl HBSS and isolated by fluorescent activated cell sorting (FACS) using a BD FACS-Aria Diva. 2000 cells per biological replicate were collected, centrifuged at 200xg for 5 mins at 4°C, washed with PBS, and resuspended in lysis buffer. RNA was extracted using Ambion RNAqueous Micro Total RNA isolation kit (AM1931, ThermoFisher Scientific). RNA integrity was checked using Bioanalyser with Agilent RNA 6000 pico kit (Agilent Technologies, 5067-1513); samples with RIN >7 were processed for library preparation. Sequencing libraries were prepared using Nextera Sample low input kit (Illumina, 15028212) and sequenced using 75bp paired end reads on the Illumina Hiseq4000 platform. A minimum of three biological replicates were used for analysis.

### Single-cell RNA extraction, library preparation and sequencing

HH6-7 embryos were electroporated with Pax2E1-EGFP to label OEPs; cells were dissociated as described above and 288 EGFP+ cells from each ss8-9, ss11-12 and ss14-15 were collected by FACS in 96-well plates. Sequencing libraries were prepared following the SmartSeq2 protocol (67). Libraries were sequenced on the Illumina NextSeq500 platform using single end 75bp sequencing (ss8-9) or on the HiSeq4000 platform using paired end 75bp sequencing (ss11-12, ss14-15).

### Nuclei isolation, ATAC library preparation and sequencing

The ATACseq library was prepared following published protocols (68). Approximately 30 pieces of the OEP territory were dissected from ss8-9 embryos, dissociated with Dounce homogenizer (tight pestle) in lysis buffer (10mM Tris-HCl, pH7.4, 10mM NaCl, 3mM MgCl2, 0.1% IGEPAL CA-630). Nuclei were pelleted at 4000 rpm at 4°C for 10 min. After removing the lysis buffer, 1.25μl Tn5 transposase (Illumina, FC-131-1024) were added to 25μl reaction volume and incubated at 37°C for 10 mins. Tagmented DNA was then purified with Mini Elute PCR purification kit (Qiagen, 28004) followed by 9 cycles of PCR enrichment using NEB High fidelity PCR kit (NEB, M0541S). The quality of ATAC libraries was assessed with Agilent Bioanalyzer with DNA High Sensitivity kit (Agilent Technologies, 5067-4627) and quantified with Kapa NGS library quantification kit (Kapa Biosystems, KK4824). The libraries were sequenced with Illumina HiSeq 2000/2500 in 2×100 cycles.

### H3K27ac, H3K27me3 and H3K4me4 ChIP library preparation and sequencing

Approximately 200 pieces of OEP ectoderm were dissected from ss8-9 embryos. The tissue was dissociated in nuclei expulsion buffer (0.5% NP-40, 0.25% Triton X-100, 10 mM Tris-HCl, pH 7.5, 3mM CaCl2, 0.25M sucrose, 1x protease inhibitor (Roche), 1mM DTT, and 0.2mM PMSF) using a Dounce homogenizer (loose pestle); crosslinking was performed in 1% formaldehyde for 9 minutes, followed by quenching with 1M glycine for 5 minutes. Cells were then pelleted and washed 3 times with PBS containing protease inhibitor (Roche, 11873580001). The pellets were snap-frozen and stored at -80°C for later use. Crosslinked chromatin was fragmented by sonication in an ice bath (Misonix Q700 at 7 Amplitude, 5×40 seconds, 30 seconds on, 60 seconds off) and immunoprecipitated following the Nano-ChIP protocol (69). For chromatin immunoprecipitation (ChIP), the following antibodies were used anti-H3K27Ac (Abcam, ab4729), anti-H3K4me3 (Diagenode, A5051-001P) and anti-H3K27me3 (Millipore, 07449). Adaptors and primers from NEBNext library preparation kit (Illumina, E6040S) were used to prepare the library following NanoChip protocol (69). The library was enriched using 14 PCR cycles and quantified with Kapa NGS library quantification kit (Kapa Biosystems, KK4824) before pooling at a concentration of 20nM. Library quality was assessed with Agilent Bioanalyzer with DNA High Sensitivity kit (Agilent Technologies, 5067-4627) and quantified with Kapa NGS library quantification kit (Kapa Biosystems, KK4824). The libraries were sequenced with Illumina HiSeq 2000/2500 in 2×100 cycles.

### High-Throughput-Sequencing-Data (HTSD) Analysis

All data alignment and downstream analysis was carried out using NF-core and custom Nextflow pipelines to allow full reproducibility. All code used, including Nextflow pipelines and downstream analysis, can be found at https://github.com/alexthiery/otic-reprogramming. Full detailed documentation for the pipeline is also available at https://alexthiery.github.io/otic-reprogramming/. A custom Docker container used for the downstream analysis pipeline can be found at https://hub.docker.com/repository/docker/alexthiery/otic-reprogramming-r_analysis. This also allows for interactive exploration of the data.

### Bulk-RNA-seq

Bulk-RNA-seq data were processed and aligned to GalGal6 using the default NF-core RNA-seq (v2.0) pipeline (70) which uses the STAR aligner. Downstream differential expression analysis (Lmx1aE1-EGFP vs Sox3U3-EGFP; Sox8OE vs ControlOE) was carried out with the DESeq2 package in R (71). Adjusted p-values were calculated using the default DESeq2 multiple test correction (Benjamini-Hochberg). Differentially expressed transcripts were determined by an absolute log2 fold-change >1.5 and adjusted p-value < 0.05.

### Single-cell-RNA-seq alignment

SmartSeq2 single-cell-RNA-seq reads were aligned and processed using a custom Nextflow DSL2 pipeline. Adaptor sequences were trimmed using Cutadapt (v2.10). HISAT2 (v2.2.1) was then used to build a genome index from GalGal6 (amended to include a GFP sequence), before extracting splice sites from the GTF and aligning reads. Read counts were obtained using HTSeq (v0.12.4). BAM files from HISAT2 were also passed to Velocyto (v0.17) (27) in order to get spliced and unspliced expression matrices for further downstream analysis.

### Single-cell-RNA-seq data analysis

Downstream data analysis was carried out primarily using the Antler R package (version: Development2019) (72). We excluded from the dataset: cells which expressed fewer than 1k genes or fewer than 500k reads, cells with more than 6% reads from mitochondrial genes, genes which were expressed in fewer than 3 cells and genes with CPM < 10.

### Identification of gene modules

To identify clusters of genes with correlated expression unbiasedly (gene modules), we used the Antler identifyGeneModules function. Genes which did not have a Spearman correlation greater than 0.3 with at least 3 other genes were first removed. Genes were then iteratively hierarchically clustered into gene modules and filtered. Gene modules were filtered based on the minimum expression level (5 CPM) and the proportion (0.4) of cells expressing a gene module. The number of final gene modules was set to 40 to achieve reasonably large gene modules, which broadly characterize cell type diversity across the dataset. These gene modules were then filtered based on the presence of genes with known expression profiles in ectodermal derivatives. Cells were then re-clustered based on the remaining gene modules, with the expression of known markers used to assign cell states. Placodal cells were then subset and re-clustered using a new set of gene modules identified from the subset cell population. Three main cell states were identified as otic, epibranchial and OEP populations.

### Gene expression dynamics at the otic-epibranchial branching point

To model the transcriptional dynamics at the otic-epibranchial branching point, we ordered cells along pseudotime using Monocle2 (73). The lineage tree was rooted to the earliest cell state (ss8-9) which was comprised mostly of OEPs. The expression of genes along the bifurcation point was then modelled using branch expression analysis modelling (BEAM).

To assess whether individual OEPs simultaneously express markers associated with the otic or epibranchial state, we calculated the co-expression of key otic and epibranchial genes within each of the Monocle branches. First, gene expression was binarized based on the presence or absence of gene expression; then the proportion of cells co-expressing pairs of otic-epibranchial genes within each of the branches was calculated. A Kruskal Wallis test was then used to compare the proportion of cells co-expressing otic and epibranchial genes between the three branches. Post-hoc pairwise comparisons between the OEP branch and the otic or epibranchial branch were carried out using two-tailed Wilcoxon rank sum tests.

RNA velocity provides an alternative to Monocle2 for inferring future cell state of individual cells. Spliced, un-spliced and spanning matrices obtained from Velocyto (27) were subset in R based on genes and cells used to generate the Monocle trajectory. RNA velocity was then calculated using the Velocyto.R package, where spanning reads were used to fit gene offsets. Single cell velocities as well as vector fields were subsequently visualized on both tSNE and Monocle DDRTree embeddings.

### ChIP-seq and ATAC-seq alignment and peak calling

ChIP-seq and ATAC-seq data were processed and aligned to GalGal6 using the default NF-core ChIP-seq (v1.2.0) and NF-core ATAC-seq (v1.2.0) pipelines (70), respectively. Reads were aligned using the Burrows-Wheeler aligner. MACS v2.2.7.1 was used to call broad peaks (FDR < 0.1) and narrow peaks (FDR < 0.05) for ChIP-seq and ATAC-seq data, respectively.

### Enhancer discovery

Putative enhancers were identified using a custom Nextflow DSL2 pipeline. First, bedtools (v2.29.2) was used to subset ATAC-seq peaks which overlap with a H3K27ac peak, whilst removing those overlapping with H3K27me3 peaks. The remaining ATAC-seq peaks were then annotated with Homer (v4.11), using a GalGal6 GTF which was filtered for protein coding genes. Promoter and exonic peaks were then removed. Motif enrichment and functional enrichment analysis were carried out using Homer and g:Profiler, respectively.

### Transcription Factor Binding Site Prediction

Enhancers that showed EGFP reporter activity in the otic placode where scanned for transcription factor binding sites (TFBS) using the RSAT-matrix-scan tool which scans DNA sequences with position-specific scoring matrices (http://rsat.sb-roscoff.fr/matrix-scan_form.cgi). (Citation: Jean Valéry Turatsinze, Morgane Thomas-Chollier, Matthieu Defrance and Jacques van Helden (2008). Using RSAT to scan genome sequences for transcription factor binding sites and cis-regulatory modules. Nat Protoc, 3, 1578-1588. Pubmed 18802439). A curated list of matrices was used (Supplementary Data 4). Both strands were scanned specifying the background model estimation for organism-specfic (Gallus gallus) and asking to return motifs with a p-value < 0.001.

## Data and materials availability

All material will be made available upon request after appropriate material transfer agreements. All data have been deposited in Gene Expression Omnibus (GEO; Accession number GSE168089). Nextflow pipelines and downstream analysis, can be found at https://github.com/alexthiery/otic-reprogramming. Full detailed documentation is available at https://alexthiery.github.io/otic-reprogramming/. A custom Docker container used can be found at https://hub.docker.com/repository/docker/alexthiery/otic-reprogramming-r_analysis.

## Acknowledgements

We thank Tatjana Sauka-Spengler and Ruth Williams for assistance with establishing ChIPseq and scRNAseq protocols, Monica Tambalo for help with tissue collection, Rosalinda Guerra, Ewa Kolano-Merlin and Chantal Hubens for excellent technical assistance, and James Briscoe for advice on single cell transcriptomics. Thanks to Claudio D. Stern for comments on the manuscript and the Streit group for discussions.

