## Supplemental Figures for "Sox8 remodels the cranial ectoderm to generate the ear"

Fig. S1.

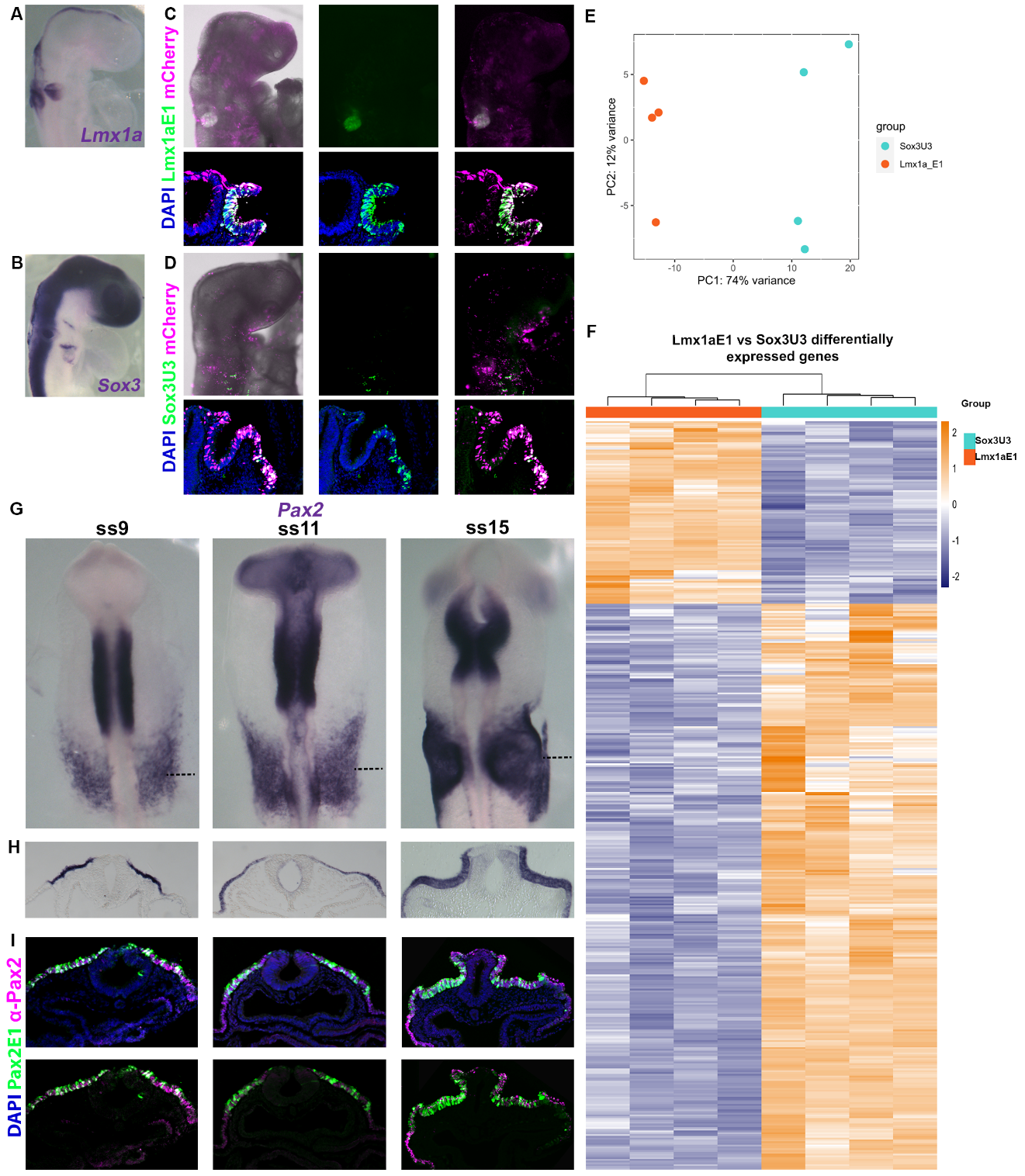

Figure S1. Transcriptomic characterisation of committed otic and epibranchial placodes. Expression of the otic marker *Lmx1a* (A) and the epibranchial marker *Sox3* (B). (C, D) Co-electroporation of EGFP reporter constructs with ubiquitously expressed mCherry (magenta) reveals enhancer activity (green), and cross sections at the level of the invaginating otic placode. Lmx1aE1-EGFP shows activity in the otic placode (C), and Sox3U3-EGFP reporter is evidenced in the epibranchial territory of ss18-20 embryos (D). (E), Principal component analysis of the RNAseq for four independent biological replicates of Lmx1aE1-EGFP+ and Sox3U3-EGFP+cells. (F), Heatmap illustrating genes differentially expressed (absolute log2 fold-change >1.5 and adjusted p-value < 0.05) between otic and epibranchial lineages. (G), *Pax2* is expressed in OEPs, otic and epibranchial cells. (H), cross section at the level of the dotted lines in (G). (I) Immunohistochemistry shows co-localisation of the Pax2E1-EGFP reporter and Pax2 endogenous protein in cross sections of embryos depicted in Fig. 1F; sections are at the same level as in G.

Fig. S2.

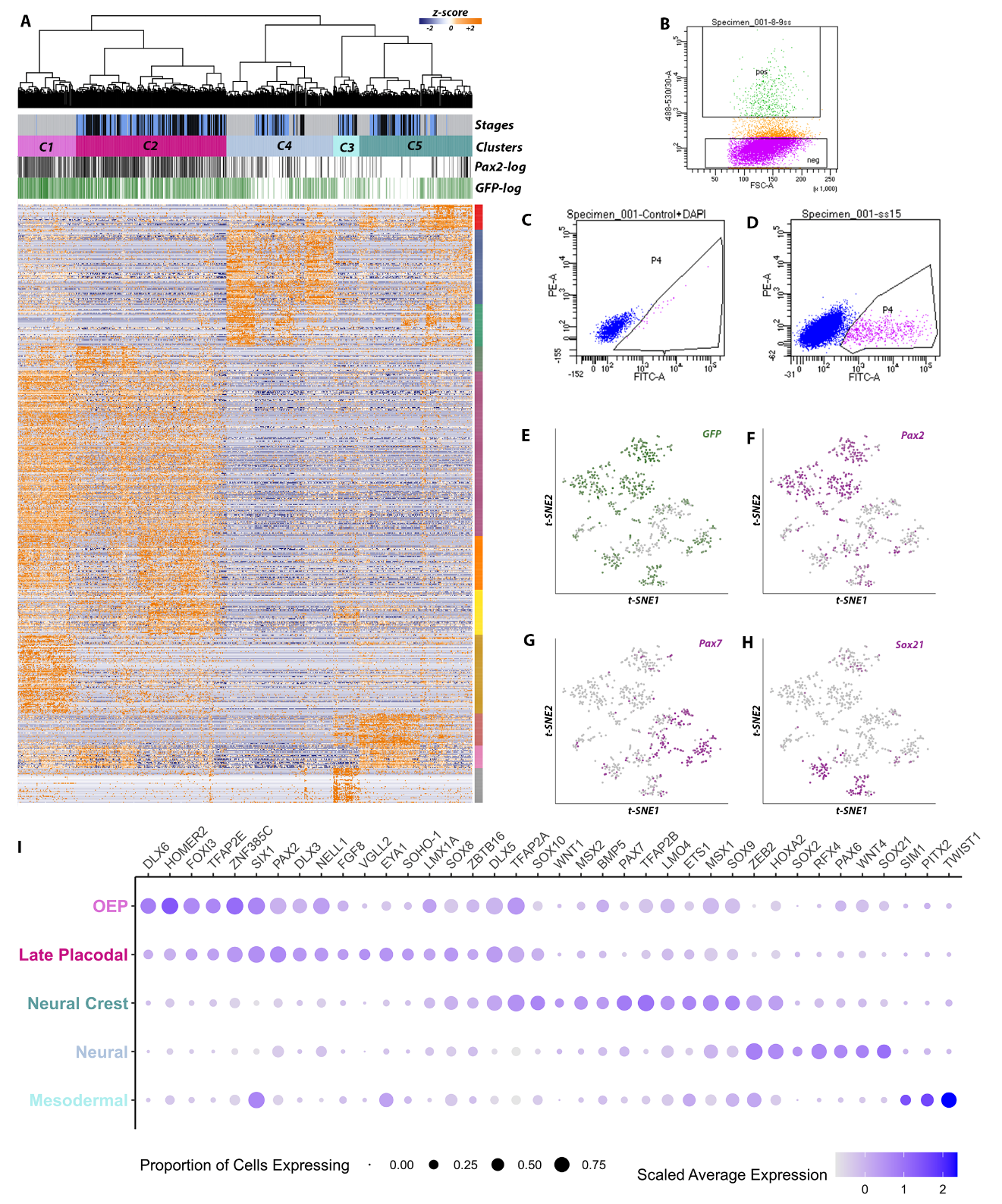

Figure S2. Hierarchical clustering of single cells from different developmental stages identifies distinct cell states. (A), Unsupervised hierarchical clustering based on the association of gene modules groups cells into five clusters C1-5. (B-D) FACS gating plots showing the selection of EGFP+ cells for ss8-9 (green positive ‘pos’ group) (B), ss11-12 (P4) (C) and ss14-15 (P4) (D) samples. (E-H), tSNE representation showing the expression of (E) *GFP* mRNA, (F) the otic-epibranchial marker *Pax2*, (G) the neural crest marker *Pax7* and (H) the neural marker *Sox21*. Note that these plots relate to Fig. 1G, H. (I), Dot plot for OEP, late placodal, neural crest, neural and mesodermal markers based on scRNAseq data showing that cell states can be broadly characterized based on combinatorial gene expression. Expression level is indicated by color intensity and gene expression frequency by the dot size.

**Fig. S3**

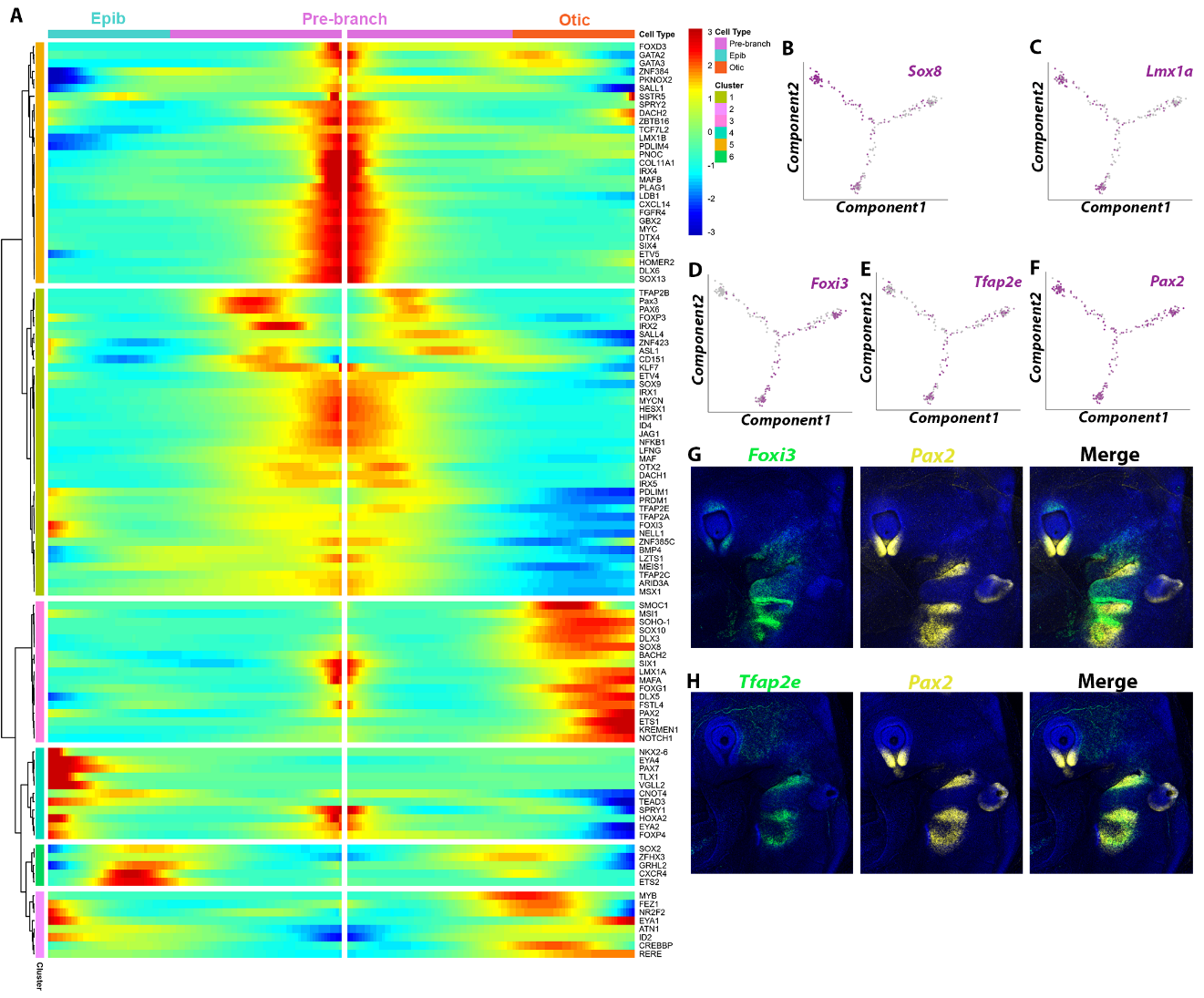

**Figure S3.** **Trajectory based expression analysis identifies core TFs of ear development.** (**A**), BEAM (Branch Expression Analysis Modelling) plot of genes in the placodal subset of cells. (**B**-**F**), Expression of OEP, otic and epibranchial markers along the pseudo-time axis as determined by Monocle2. (**G**, **H**) HCR at HH17 shows that *Foxi3* (**G**) and *Tfap2e* (**H**) co-localize with the well-known placodal marker *Pax2* and are expressed in the epibranchial placodes.

Fig. S4

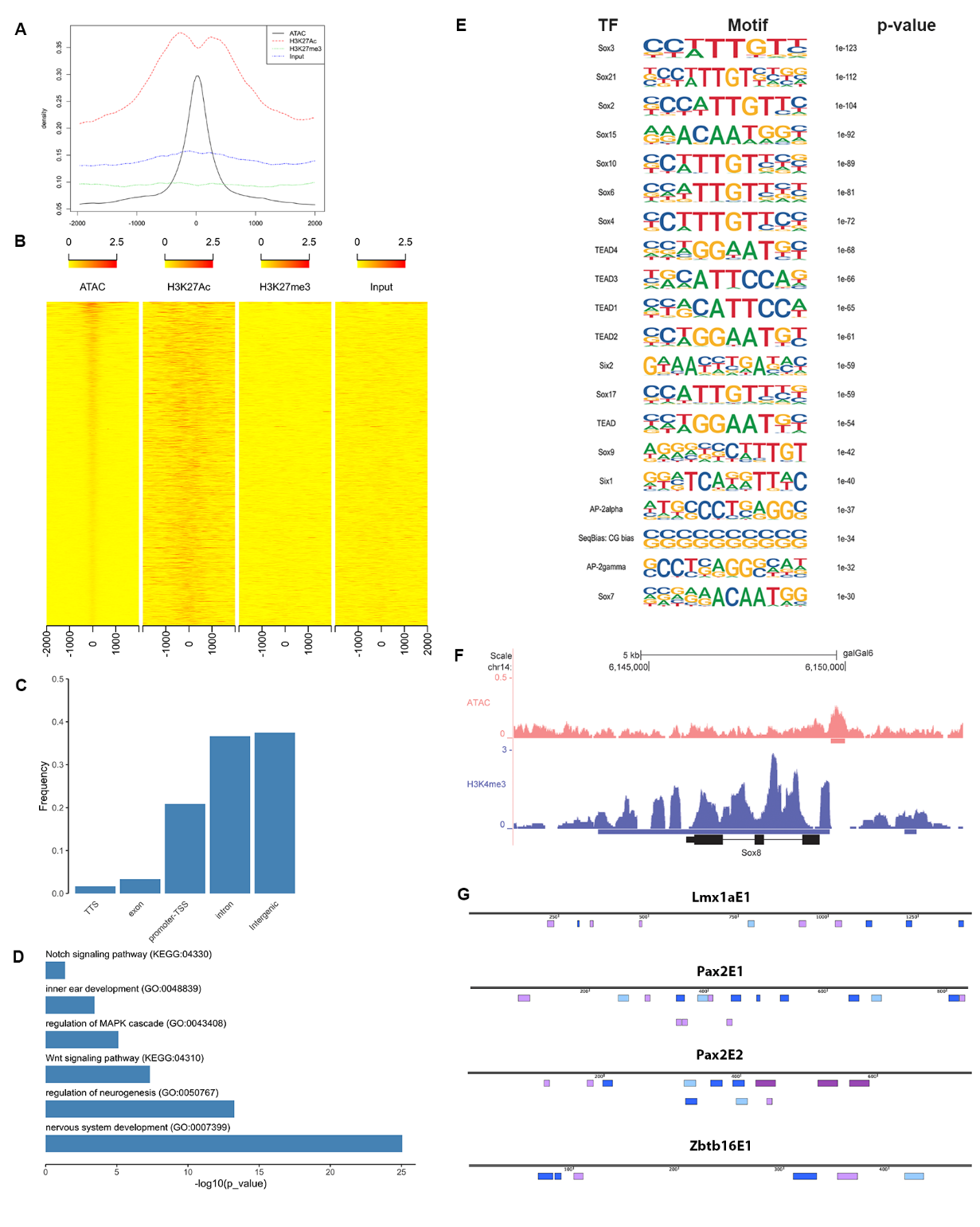

**Figure S4.** **Profiling of chromatin accessibility in OEPs.** (**A**) Plot showing the distribution profiles for ATACseq, ChIPseq for H3K27ac and for H3K27me3 and input. This shows bimodal distribution of H3K27ac surrounding ATAC peaks. (**B**) Heatmap showing the localization of ATACseq, ChIPseq for H3K27ac, H3K27me3 and input peaks. (**C**) Annotation of ATACseq peaks according to genomic location. (**D**) Functional enrichment analysis of predicted enhancers shows enrichment of Gene Ontology (GO) terms related to early ear development. (**E**) Transcription factor motif enrichment analysis of ear enhancers reveals binding sites for the Sox, Tead, Six and TFAP families as highly enriched. (**F**) The Sox8 locus is decorated with broad H3K4m3 peaks. (**G**) Transcription factor binding site prediction using RSAT shows predicted motifs for the core-components of the ear-network Sox8 (blue), Pax2 (lilac), Lmx1a (light-blue) and Zbtb16 (purple) in the enhancers associated with Lmx1a, Pax2 and Zbtb16 loci.

Fig. S5

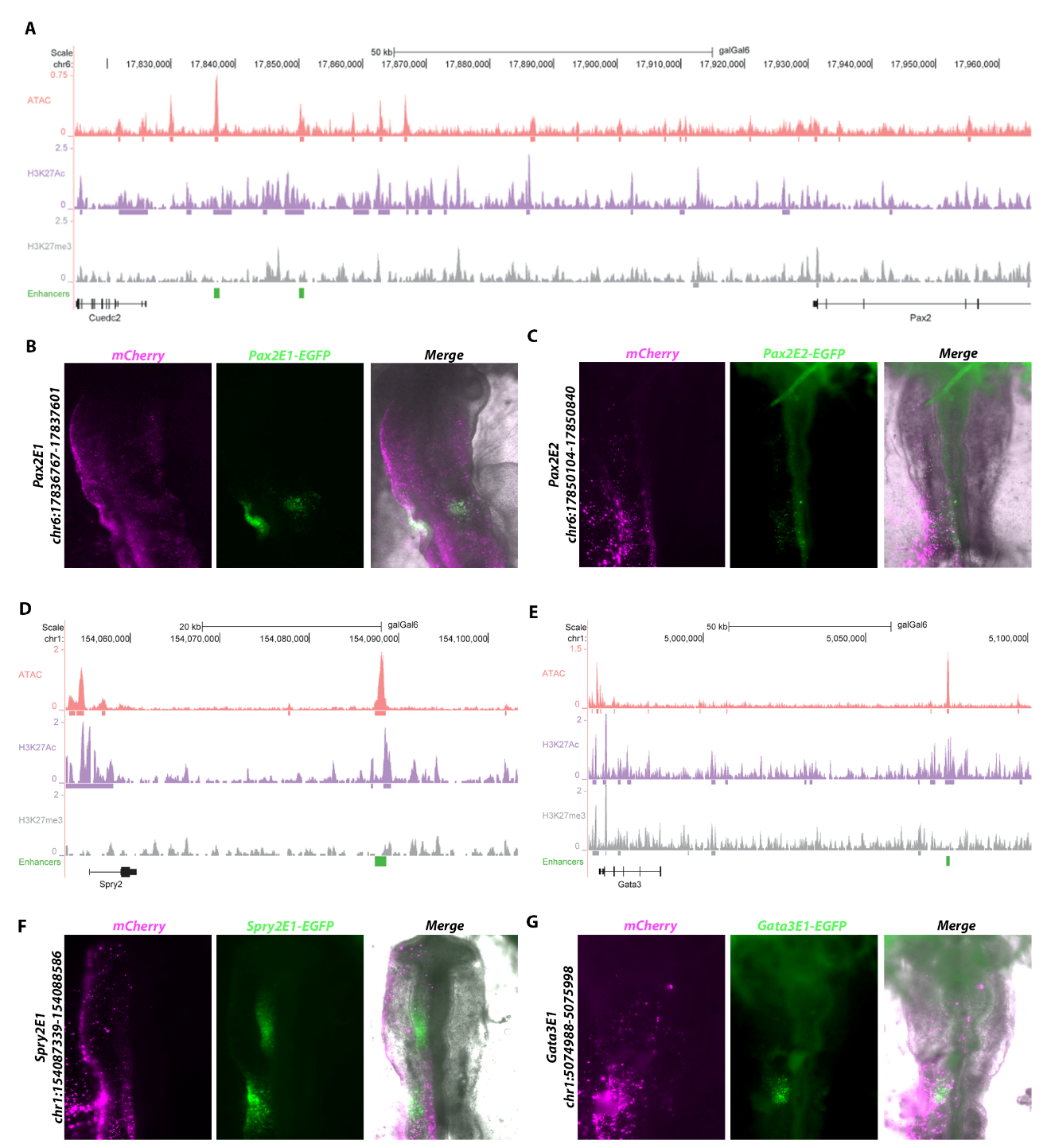

**Figure S5**. **Chromatin accessibility and ChIPseq identifies new otic enhancers.** (**A**), (**D**), (**E**), ATACseq, H3K27ac and H3K27me3 ChIPseq tracks surrounding the Pax2, Spry2 and Gata3 loci. Regions labelled in green were cloned into EGFP reporter constructs. (**B**), (**C**), (**F**), (**G**), Co-electroporation of the EGFP reporter constructs with ubiquitously expressed mCherry (magenta) reveals enhancer activity (green) in the otic-epibranchial territory for Pax2E1EGFP (ss13) and Pax2E2-EGFP (ss9) (**B**, **C**) in OEPs and the mid-brain-hindbrain boundary for Spry2-EGFP (ss10) (**F**) and the otic placode for Gata3E1-EGFP (ss12) (**G**).

Fig. S6

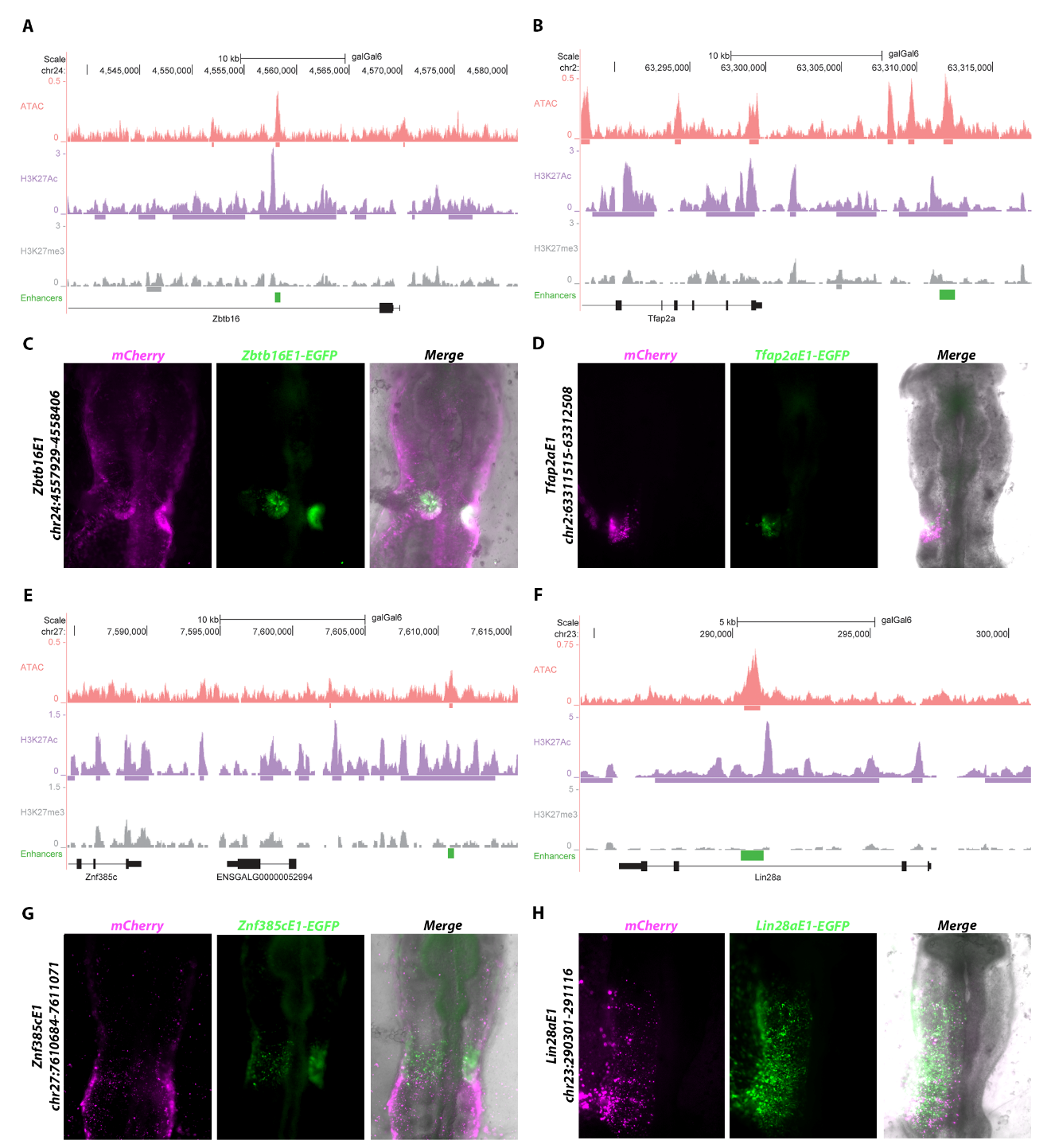

**Figure S6.** **Chromatin accessibility and ChIPseq identifies new otic enhancers.** (**A**), (**B**), (**E**), (**F**), ATACseq, H3K27ac and H3K27me3 ChIPseq tracks surrounding the Zbtb16, Tfap2a, Znf385c and Lin28a loci. Regions labelled in green were cloned into EGFP reporter constructs. (**C**), (**D**), (**G**), (**H**), Co-electroporation of the EGFP reporter constructs with ubiquitously expressed mCherry (magenta) reveals enhancer activity (green) in the otic placode for Zbtb16E1-EGFP (ss13) (**C**) and for TFAP2aE1-EGFP (ss11) (**D**), in OEPs for Znf385c-EGFP (ss12) (**G**) and the cranial ectoderm including OEPs for Lin28aE1-EGFP (ss10) (**H**).

**Fig. S7**

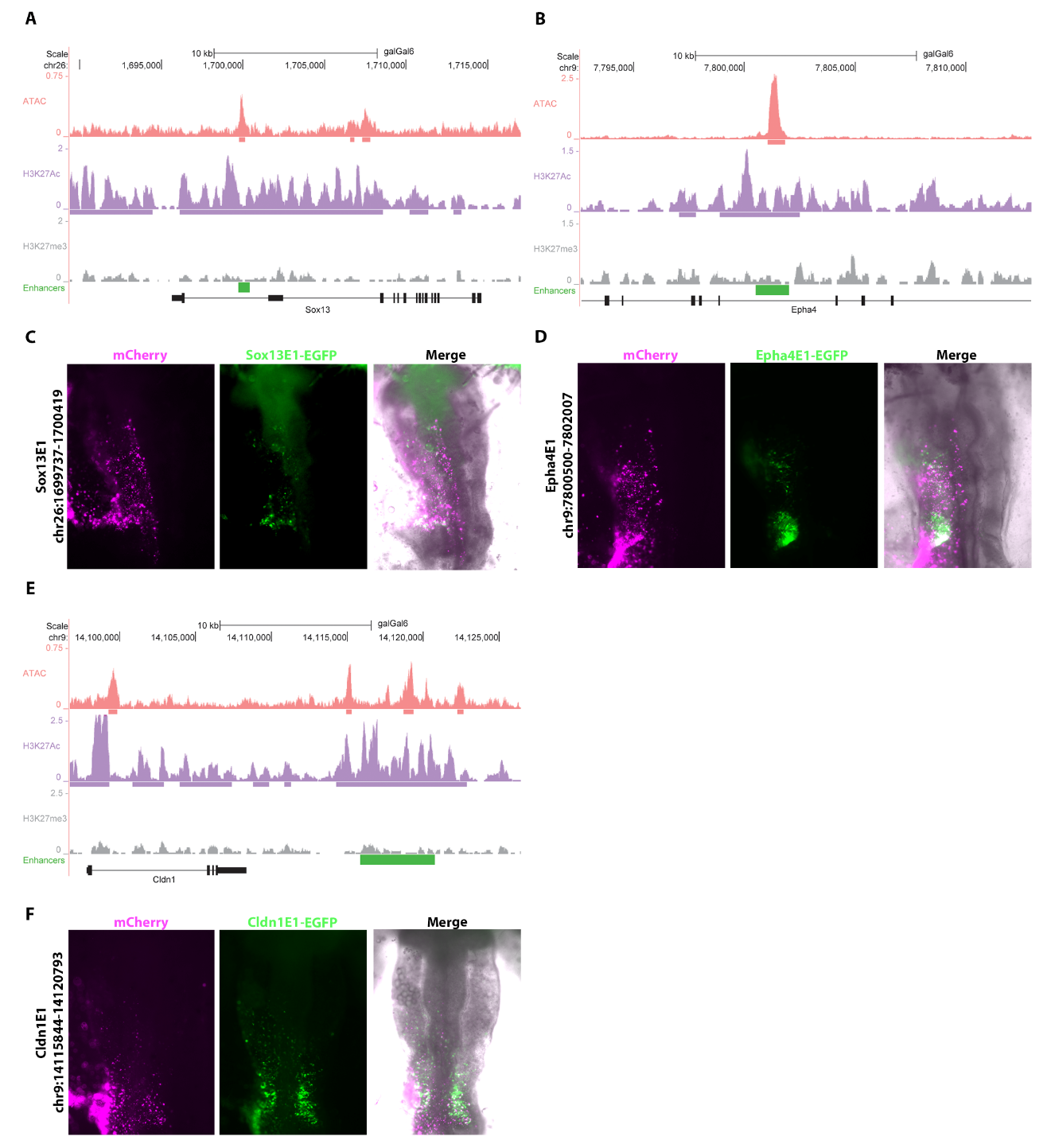

**Fig. S7. Chromatin accessibility and ChIPseq identifies new otic enhancers.** (**A**), (**B**), (**E**), ATACseq, H3K27ac and H3K27me3 ChIPseq tracks surrounding the Sox13, EphA4 and Cldn1 loci. Regions labelled in green were cloned into EGFP reporter constructs. (**C**), (**D**), (**F**), Co-electroporation of the EGFP reporter constructs with ubiquitously expressed mCherry (magenta) reveals enhancer activity (green) in OEPs for Sox13-EGFP (ss12) (**C**), in the otic placode for EphA4E1-EGFP (ss13) (**D**) and in OEPs for Cldn1-EGFP (ss11) (**G**). All three show variable activity in some cells of the cranial ectoderm.

Fig. S8

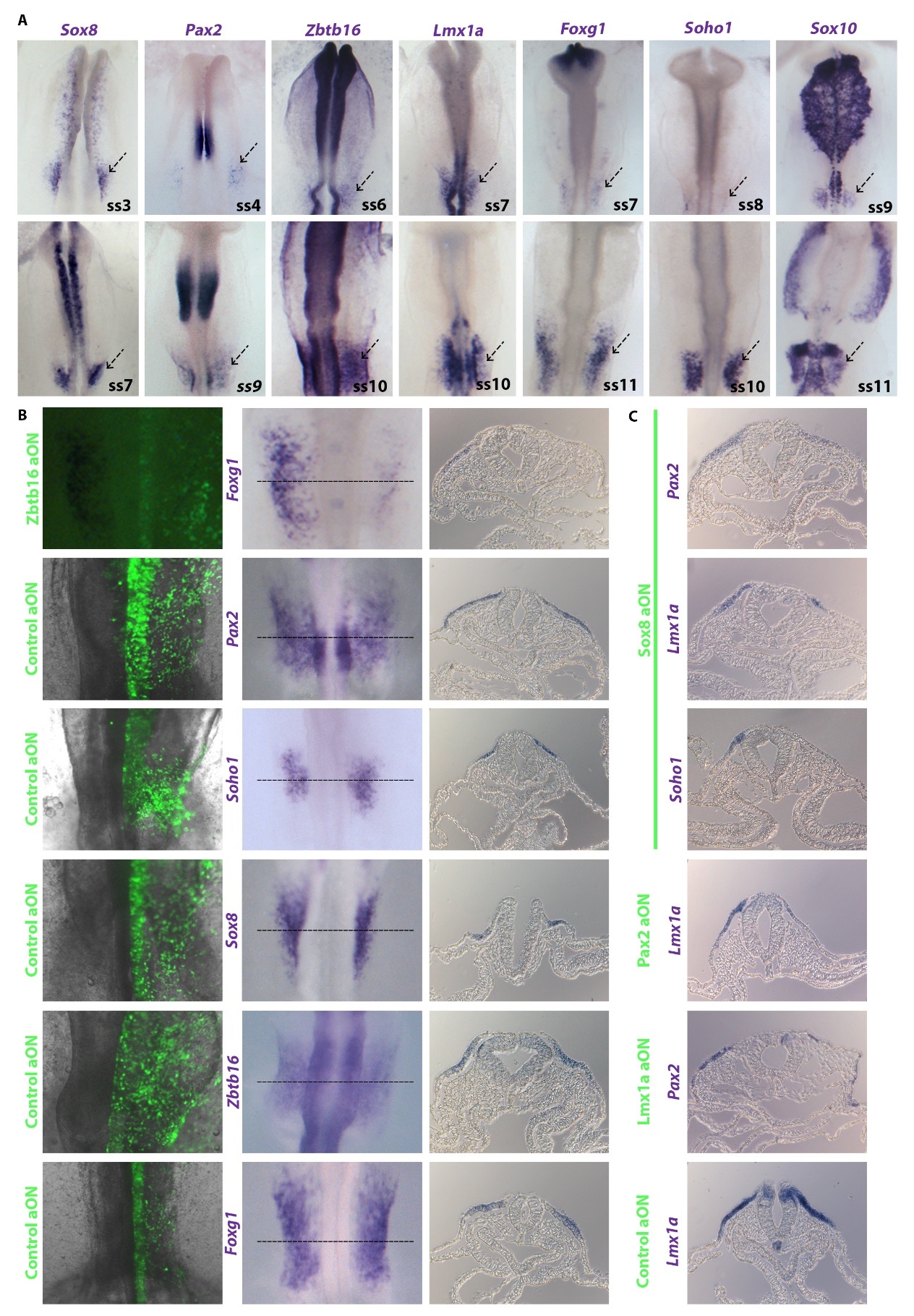

**Fig. S8. Hierarchy of expression and regulatory relationship of otic enriched transcription factors.** (**A**), *In situ* hybridization shows that *Sox8* is the earliest transcription factor tested to be expressed in OEPs (ss3), followed by *Pax2* (ss4), *Zbtb16* (ss6) and *Lmx1a* (ss7). *Foxg1* is first expressed weakly at ss7, while *Soho1* (ss8) and *Sox10* (ss9) begin to be expressed as the otic placode becomes visible. Arrows indicate the OEP/otic placode territory. (**B**), Knock-down using antisense oligonucleotides for Zbtb16 and control antisense oligonucleotides followed by *in situ* hybridization. Knock-down of Zbtb16 results in downregulation of *Foxg1*, while controls have no effect as observed in both whole mounts and sections at the level of the otic placode (dashed line). (**C**) Transverse sections at the level of the otic territory of embryos depicted in Figure 3E show the reduced marker gene expression on the aON targeted side (right), but not on the contralateral uninjected side (left). Control aONs have do not have any effect.

**Fig. S9**

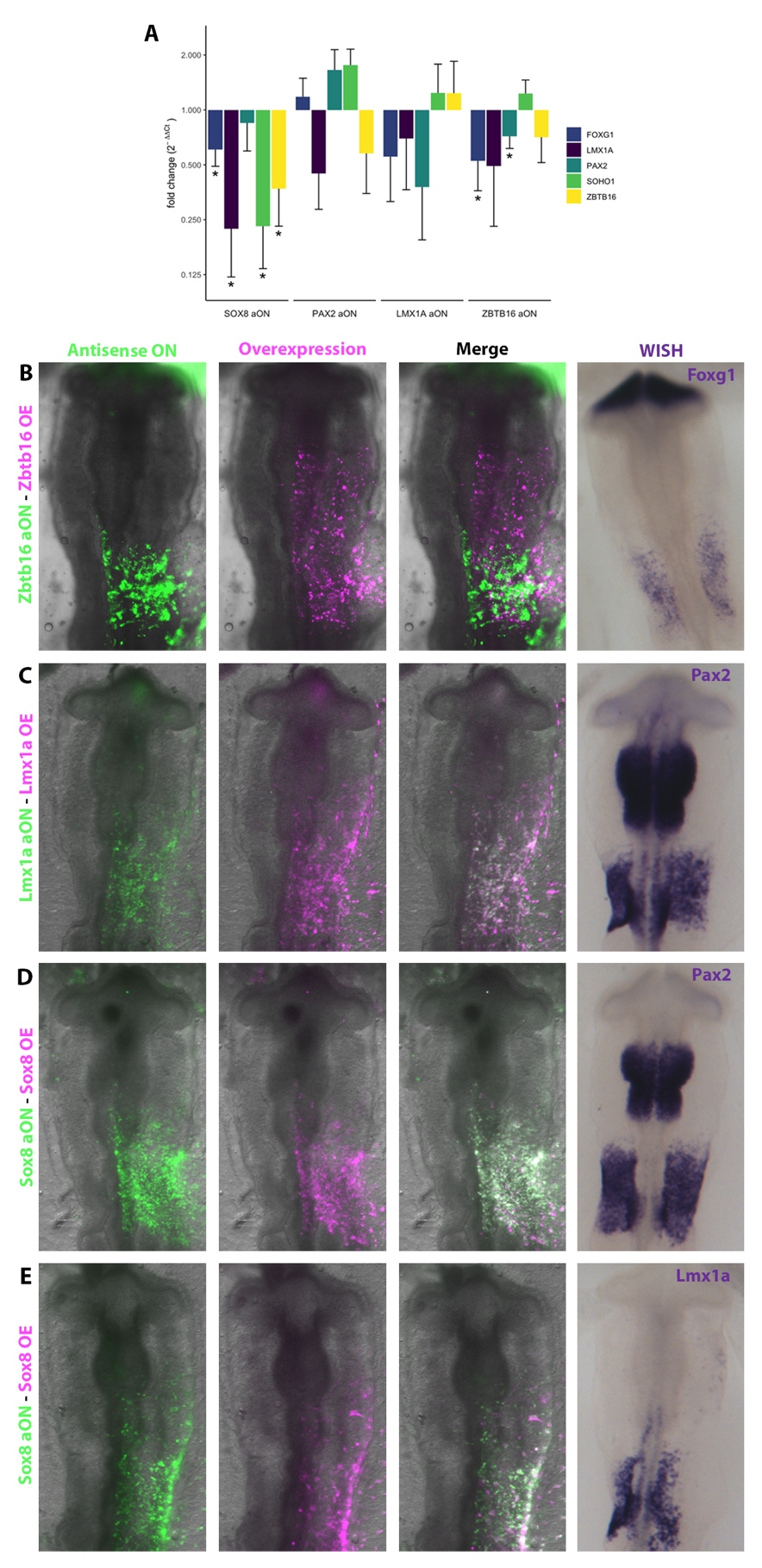

**Fig. S9. Regulatory relationship of otic enriched transcription factors and rescue by overexpression of the depleted factors. (A)** Knock-down using antisense oligonucleotides followed by qPCR analysis shows that Sox8 is required for the expression of all transcription factors tested, while Pax2 is necessary for *Lmx1a*, Lmx1a for *Pax2*, and Zbtb16 for *Foxg1* and *Pax2* expression. Bars represent indicate fold change (2-ΔΔCt) relative to controls. Fold change plotted on log2 scale for biological interpretability. Error bars indicate ±1 standard deviation. * p-value < 0.05.  **(B-E)** Co-electroporation of antisense oligonucleotides (aON) green) and the construct harboring the full-length sequence of the corresponding gene (magenta) shows rescue of *Foxg1* expression for Zbtb16 aON (**B**), *Pax2* for Lmx1a aON (**C**), *Pax2* and *Lmx1a* for Sox8 aON (**D, E**).

**Fig. S10**

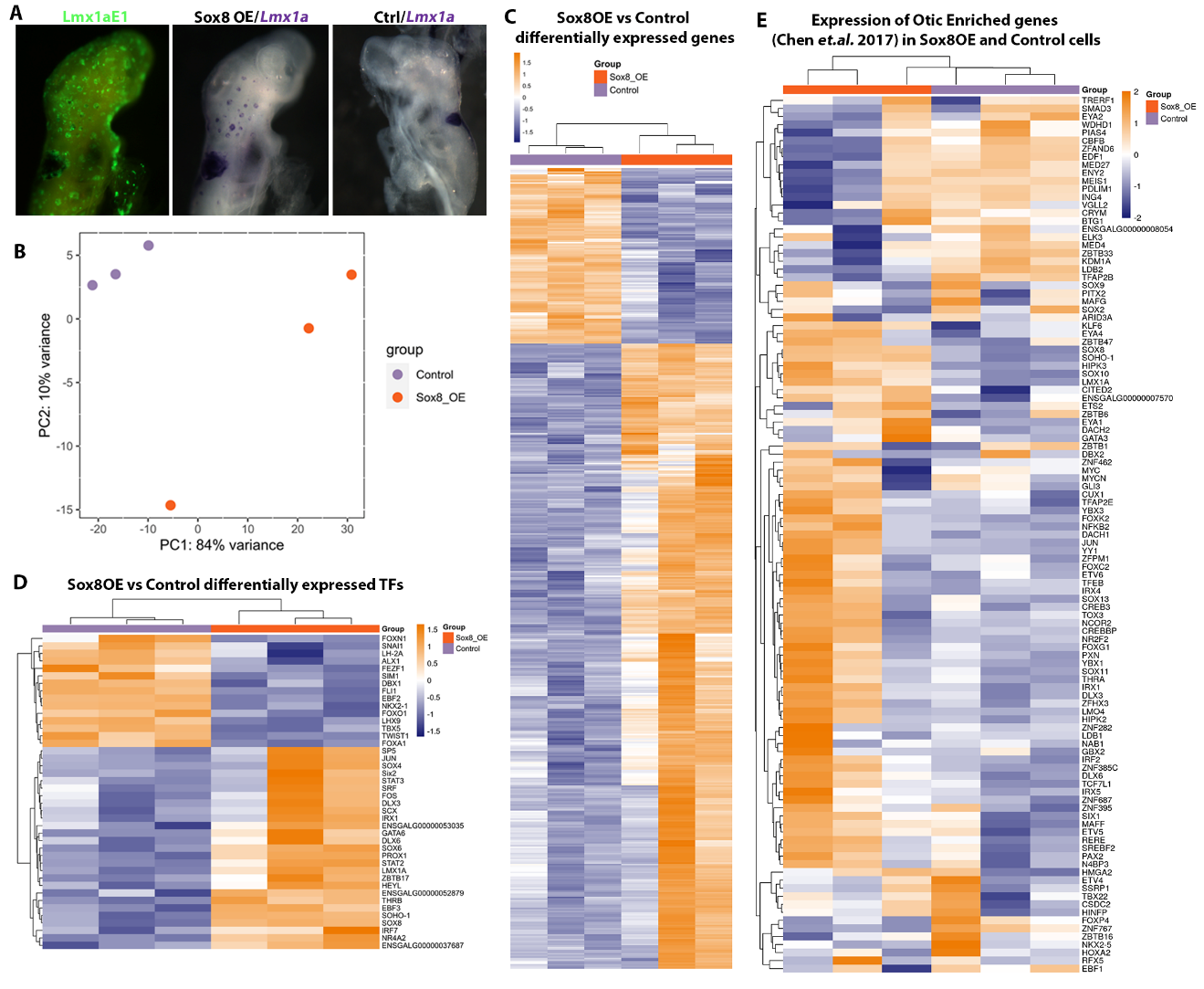

**Fig. S10. Sox8 overexpression in the cranial ectoderm activates the otic programme.** (**A**), Overexpression of Sox8 triggers the activation of Lmx1aE1-EGFP (green) and Lmx1a transcript in ectopic vesicles as compared to the non-electroporated contralateral side. (**B**) Principal component analysis of three independent biological replicates for RNAseq of control and Sox8 overexpressing cells. (**C**) Heatmap of differently expressed TFs in the double positive Sox8-overexpressing/Lmx1aE1+ cells shows upregulation of otic genes compared to controls (absolute log2 fold-change >1.5 and adjusted p-value < 0.05). (**D**) Heatmap of differentially expressed genes between Sox8OE/Lmx1aE1+ cells versus mCherry/EGFP double positive control cells (absolute log2 fold-change >1.5 and adjusted p-value < 0.05). (**E**) Heatmap showing the expression of otic enriched genes from Chen *et.al.* 2017 (45) within the Sox8OE DE genes. Of 110 genes, 98 (89%) are upregulated in at least one sample of Sox8OE-derived cells but not in controls.

**Dataset S1 (separate file).**

Supplementary Table 1: This table shows the differential expression results for genes with absolute log2FC > 1.5 and adjusted p-value < 0.05 when comparing Lmx1a_E1 and Sox3U3 samples (Lmx1a_E1 - Sox3U3).

**Dataset S2 (separate file).**

Supplementary Table 2: This table shows the gene modules (GM) that characterise OEP, Otic and Epibranchial cell states.

**Dataset S3 (separate file).**

Supplementary Table 3: This file shows the predicted Transcription Factor Binding Sites (TFBS) of the enhancers actve in the active placode shown in this paper (Fig2A, Fig. S5-7). Analysis was performed as described in the Methods section using RSAT tool (<http://rsat.sb-roscoff.fr/matrix-scan_form.cgi>).

**Dataset S4 (separate file).**

Supplementary file 4: This file contains the position-specific scoring matrices (PSSM) used to perform the TFBS prediction on RSAT online tool as described in the Methods section.

**Dataset S5 (separate file).**

Supplementary Table 5: “DE genes in Sox8OE vs Control” shows the differential expression results for genes with absolute log2FC > 1.5 and adjusted p-value < 0.05 when comparing Sox8 overexpression and control samples.

**Dataset S6 (separate file).**

Supplementary Table 6: This table shows differentially expressed (absolute FC > 1.5 and padj (FDR) < 0.05) transcription factors between Sox8 overexpression and control samples (Sox8 - Control).

**Dataset S7 (separate file).**

Supplementary Table 7: This table shows the experiments performed using different combinations of TFs to test their ability to generate ectopic otic vesicles in the head ectoderm. Experiments were performed as described in the Methods section for Sox8 over expression electroporation.
